# Site-specific N-glycosylation Characterization of Recombinant SARS-CoV-2 Spike Proteins

**DOI:** 10.1101/2020.03.28.013276

**Authors:** Yong Zhang, Wanjun Zhao, Yonghong Mao, Yaohui Chen, Shisheng Wang, Yi Zhong, Tao Su, Meng Gong, Dan Du, Xiaofeng Lu, Jingqiu Cheng, Hao Yang

## Abstract

The glycoprotein spike (S) on the surface of SARS-CoV-2 is a determinant for viral invasion and host immune response. Herein, we characterized the site-specific N-glycosylation of S protein at the level of intact glycopeptides. All 22 potential N-glycosites were identified in the S-protein protomer and were found to be preserved among the 753 SARS-CoV-2 genome sequences. The glycosites exhibited glycoform heterogeneity as expected for a human cell-expressed protein subunits. We identified masses that correspond to 157 N-glycans, primarily of the complex type. In contrast, the insect cell-expressed S protein contained 38 N-glycans, primarily of the high-mannose type. Our results revealed that the glycan types were highly determined by the differential processing of N-glycans among human and insect cells. This N-glycosylation landscape and the differential N-glycan patterns among distinct host cells are expected to shed light on the infection mechanism and present a positive view for the development of vaccines and targeted drugs.

## INTERODUCTION

The spread of a novel severe acute respiratory syndrome coronavirus (SARS-CoV-2) has caused a pandemic of coronavirus disease 2019 (COVID-19) worldwide. Distinguished from severe acute respiratory syndrome coronavirus (SARS-CoV) and Middle East respiratory syndrome coronavirus (MERS-CoV), SARS-CoV-2 transmits more rapidly and efficiently from infected individuals, even those without symptoms, to healthy humans, frequently leading to severe or lethal respiratory symptoms(1, 2). The World Health Organization has declared the spread of SARS-CoV-2 a Public Health Emergency of International Concern. As of April 16, 2020, the virus has led to over two million confirmed cases. From SARS-CoV to SARS-CoV-2, the periodic outbreak of highly pathogenic coronavirus infections in humans urgently calls for strong prevention and intervention measures. However, there are no approved vaccines or effective antiviral drugs for either SARS-CoV or SARS-CoV-2.

Human coronaviruses, including HCoV 229E, NL63, OC43, and HKU1, are responsible for 10-30% of all upper respiratory tract infections in adults. SARS-CoV-2 can actively replicate in the throat; however, this virus predominately infects the lower respiratory tract via the envelope Spike (S) protein(2, 3). Due to its high exposure on the viral surface, the S protein can prime a protective humoral and cellular immune response, thus commonly serving as the main target for antibodies, entry inhibitors and vaccines(4-6). It has been found that human sera from recovered COVID-19 patients can neutralize SARS-CoV-2 S protein-overexpressed pseudovirions effectively(7). The use of convalescent sera in the clinic is actively undergoing a comprehensive evaluation. However, the passive antibody therapy with convalescent sera would be a stopgap measure and may not provide a protective immunity owing to their limited cross reactions(7, 8). Therefore, specific neutralizing antibodies and vaccines against SARS-CoV-2 are in rapid development for providing potent and long-lasting immune protection(9-11).

A mature SARS-CoV-2 has four structural proteins, including the S protein, envelope (E) protein, membrane (M) protein, and nucleocapsid (N) protein(1). Given its indispensable role in viral entry and infectivity, the S protein is probably the most promising immunogen, especially given the comprehensive understanding of the structure and function provided by recent studies(4, 12-14). The S protein is comprised of an ectodomain, a transmembrane anchor, and a short C-terminal intracellular tail(15). The ectodomain consists of a receptor-binding S1 subunit and a membrane-fusion S2 subunit. Following attachment to the host cell surface via S1, the S protein is cleaved at multiple sites by host cellular proteases, consequently mediating membrane fusion and making way for the viral genetic materials to enter the host cell(4, 6, 16). The S protein can bind to the angiotensin-converting enzyme II (ACE2) receptor on host cells(13, 17). The recognition of the S protein to the ACE2 receptor primarily involves extensive polar residue interactions between the receptor-binding domain (RBD) and the peptidase domain of ACE2(13, 14). The S protein RBD is located in the S1 subunit and undergoes a hinge-like dynamic movement to capture the receptor through three grouped residue clusters. Consequently, the S protein of SARS-CoV-2 displays an up to 10–20-fold higher affinity for the human ACE2 receptor than that of SARS-CoV, supporting the higher transmissibility of this new virus(13, 14).

Apart from the structural information at the residue level, the trimeric S protein is highly glycosylated, possessing 22 potentially N-linked glycosylation motifs (N-X-S/T, X≠P) in each protomer(18-20). The N-glycans on the S protein play a pivotal role in proper protein folding and protein priming by host proteases. Importantly, glycosylation is an underlying mechanism for coronavirus to evade both innate and adaptive immune responses of their hosts, as the glycans might shield the amino acid residues of viral epitopes from cell and antibody recognition(4, 5, 21). Cryo-EM has revealed the N-glycosylation on 14–16 of 22 potential sites in the SARS-CoV-2 S protein protomer(4, 13). However, these glycosites and their glycan occupancies need to be experimentally identified in detail. Glycosylation analysis via glycopeptides can provide insight into the N-glycan microheterogeneity of a specific site(22). Therefore, further identification of site-specific N-glycosylation information of the SARS-CoV-2 S protein, including that regarding intact N-glycopeptides, glycosites, glycan compositions, and the site-specific number of glycans, could be meaningful to obtain a deeper understanding of the mechanism of the viral invasion and provide guidance for vaccine design and antiviral therapeutics development(4, 23)

Herein, we characterized the site-specific N-glycosylation of recombinant SARS-CoV-2 S proteins by analysis of the intact glycopeptides using tandem mass spectrometry (MS/MS). Based on an integrated method(24), we identified 22 potential N-glycosites and their corresponding N-glycans from the recombinant S protein. All of these glycosites were found to be highly conserved among SARS-CoV-2 genome sequences. The glycosite-specific occupancy by different glycoforms was resolved and compared among S protein subunits expressed in human cells and insect cells. These detailed glycosylation profiles decoded from MS/MS analysis are expected to facilitate the development of vaccines and therapeutic drugs against SARS-CoV-2.

## EXPERIMENTAL PROCEDURES

### Materials and chemicals

Dithiothreitol (DTT), iodoacetamide (IAA), formic acid (FA), trifluoroacetic acid (TFA), TRIS base, and urea were purchased from Sigma (St. Louis, MO, USA). Acetonitrile (ACN) was purchased from Merck (Darmstadt, Germany). The zwitterionic hydrophilic interaction liquid chromatography (Zic-HILIC) materials were obtained from Fresh Bioscience (Shanghai, China). Commercially available recombinant SARS-CoV-2 S protein (S1+S2 ECD and RBD, His tag) expressed in insect cells (High Five) via baculovirus and S protein (S1 and RBD, His tag) expressed in human embryonic kidney cells (HEK293) were purchased from Sino Biological (Beijing, China). Sequencing-grade trypsin and Glu-C were obtained from Enzyme & Spectrum (Beijing, China). The quantitative colorimetric peptide assay kit was purchased from Thermo Fisher Scientific (Waltham, MA, USA). Deionized water was prepared using a Milli-Q system (Millipore, Bedford, MA, USA). All other chemicals and reagents of the best available grade were purchased from Sigma-Aldrich or Thermo Fisher Scientific.

### Protein digestion

The recombinant S proteins were proteolyzed using an in-solution protease digestion protocol. In brief, 50 μg of protein in a tube was denatured for 10 min at 95 °C. After reduction by DTT (20 mM) for 45 min at 56 °C and alkylating with IAA (50 mM) for 1 h at 25 °C in the dark, 2 μg of protease (trypsin or/and Glu-C) was added to the tube and incubated for 16 h at 37 °C. After desalting using a pipette tip packed with a C18 membrane, the peptide concentration was determined using a peptide assay kit based on the absorbance measured at 480 nm. The peptide mixtures (intact N-glycopeptides before enrichment) were freeze-dried for further analysis.

### Enrichment of intact N-glycopeptides

Intact N*-*glycopeptides were enriched with Zic-HILIC (Fresh Bioscience, Shanghai, China). Specifically, 20 μg of peptides were resuspended in 100 μL of 80% ACN/0.2% TFA solution, and 2 mg of processed Zic-HILIC was added to the peptide solution and rotated for 2 h at 37 °C. Finally, the mixture was transferred to a 200-μL pipette tip packed with a C8 membrane and washed twice with 80% ACN/0.2% TFA. After enrichment, intact N*-*glycopeptides were eluted three times with 70 μL of 0.1% TFA and dried using a SpeedVac for further analysis.

### Deglycosylation

Enriched intact N*-*glycopeptides were digested using 1 U PNGase F dissolved in 50 μL of 50 mM NH_4_HCO_3_ for 2 h at 37 °C. The reaction was terminated by the addition of 0.1% formic acid (FA). The deglycosylated peptides were dried using a SpeedVac for further analysis.

### Liquid chromatography-MS/MS analysis

All samples were analyzed using SCE-higher-energy collisional dissociation (HCD)-MS/MS with an Orbitrap Fusion Lumos mass spectrometer (Thermo Fisher Scientific). In brief, intact N-glycopeptides before or after enrichment and deglycosylated peptides were dissolved in 0.1% FA and separated on a column (ReproSil-Pur C18-AQ, 1.9 μm, 75 μm inner diameter, length 20 cm; Dr Maisch) over a 78-min gradient (buffer A, 0.1% FA in water; buffer B, 0.1% FA in 80% ACN) at a flow rate of 300 nL/min. MS1 was analyzed using a scan range (m/z) of 800–2000 (intact N-glycopeptides before or after enrichment) or 350–1550 (deglycosylated peptides) at an Orbitrap resolution of 120,000. The RF lens, AGC target, maximum injection time, and exclusion duration were 30%, 2.0 e^4^, 100 ms, and 15 s, respectively. MS2 was analyzed with an isolation window (m/z) of 2 at an Orbitrap resolution of 15,000. The AGC target, maximum injection time, and the HCD type were standard, 250 ms, and 30%, respectively. The stepped collision mode was turned on with an energy difference of ±10%.

### Data Analysis

The raw data files were searched against the SARS-CoV-2 S protein sequence using Byonic software (version 3.6.0, Protein Metrics, Inc.), with the mass tolerance for precursors and fragment ions set at ±10 ppm and ±20 ppm, respectively. Two missed cleavage sites were allowed for trypsin or/and Glu-C digestion. The fixed modification was carbamidomethyl (C), and variable modifications included oxidation (M), acetyl (protein N-term), and deamidation (N). In addition, 38 insect N-glycans or 182 human N-glycans were specified as N-glycan modifications for intact N-glycopeptides before or after enrichment. We then checked the protein database options, including the decoy database. All other parameters were set at the default values, and protein groups were filtered to a 1% false discovery rate based on the number of hits obtained for searches against these databases. Stricter quality control methods for intact N-glycopeptides and peptide identification were implemented, requiring a score of no less than 200 and identification of at least six amino acids.

Furthermore, all of these peptide spectrum matches (PSMs) and glycopeptide-spectrum matches (GPSMs) were examined manually and filtered using the following standard criteria: PSMs were accepted if there were at least 3 b/y ions in the peptide backbone, and GPSMs were accepted if there were at least two glycan oxonium ions and at least 3 b/y ions in the peptide backbone. Quantitative analysis of the intact N-glycopeptide were performed as described previously(25). N-glycosite conservation analysis was performed using R software packages. Model building based on the Cryo-EM structure (PDB: 6VSB) of SARS-CoV-2 S protein was performed using PyMOL.

### Statistical analysis

The number of intact N-glycopeptides and N-glycans identified from triplicate experimental replicates was analyzed by Student’s t-test for statistical comparison between the two groups, before and after enrichment. Data were presented as means ± SD. A *P*-value < 0.05 was considered significant.

## RESULTS

### Strategy for site-specific N-glycosylation characterization

Previous studies have revealed that the glycosylated coronavirus S protein plays a critical role in the induction of neutralizing antibodies and protective immunity. However, the glycans on S protein might also surround the protein surface and form an immunologically inert “self” glycan shield for virus evasion from the immune system(5, 21, 26). Herein, we aimed to decode the detailed site-specific N-glycosylation profile of the SARS-CoV-2 S protein. The native S protein ectodomain expressed by a baculovirus expression vector in insect cells with no amino acid substitution was first used to analyze the glycosylation landscape, since this expression system can express the S protein without resulting in splicing via host proteases. Moreover, insect cells can mimic the process of mammalian cell glycosylation(4, 27). The S protein of the human coronavirus NL63 (HCoV-NL63) has been successfully expressed in insect cells for the resolution of its protein structure and glycan shield(21). The recombinant SARS-CoV-2 S ectodomain contains 1209 amino acids (residues 16–1,213) that are translated from a complete genome sequence (GenBank: MN908947.3)(28) and includes 22 putative N-glycosylation sequons (motif N-X-S/T, X≠P). Theoretical analysis of the enzymatic sites showed that trypsin alone did not produce peptides of sufficiently appropriate length of peptides to cover all potential N-glycosites (Fig. S1A). The missing potential N-glycosites were found back by introducing the endoproteinase Glu-C (Fig. S1B). Hence, we took advantage of this complementary trypsin and Glu-C digestion approach by using either a single enzyme or dual ones (Fig. S1C). Meanwhile, the recombinant SARS-CoV-2 S protein S1 subunit expressed in human cells was obtained for analysis of the site-specific N-glycans, as the N-glycan compositions in insect cells would be different from those in native human host cells(27). The S1 subunit contains 681 amino acids (residues 16–685) and 13 potential N-glycosites. Trypsin alone or dual digestion could cover all potential N-glycosites (Fig. S1C). By doing so, each N-glycosylation sequon of the different recombinant proteins was covered by glycopeptides of a suitable length for achieving good ionization and fragmentation.

To raise the abundance of intact glycopeptides, zwitterionic hydrophilic interaction liquid chromatography (Zic-HILIC) materials were used to enrich glycopeptides. Concurrently, the enrichment of intact N-glycopeptides can reduce signal suppression from unglycosylated peptides. However, there are no materials available that can capture all glycopeptides without preference. For these reasons, site-specific glycosylation was determined based on a combined analysis of the intact N-glycopeptides before and after enrichment. Furthermore, the deglycosylated peptides following enrichment were used to confirm or retrieve the N-glycosites by removing potential interferences from glycans. In brief, the integration of complementary digestion and N-glycoproteomic analysis at three levels (before and after enrichment, and at the deglycopeptides levels) is a promising approach to comprehensively and confidently profile the site-specific N-glycosylation of recombinant SARS-CoV-2 S proteins (Fig. 1).

**Figure 1.**
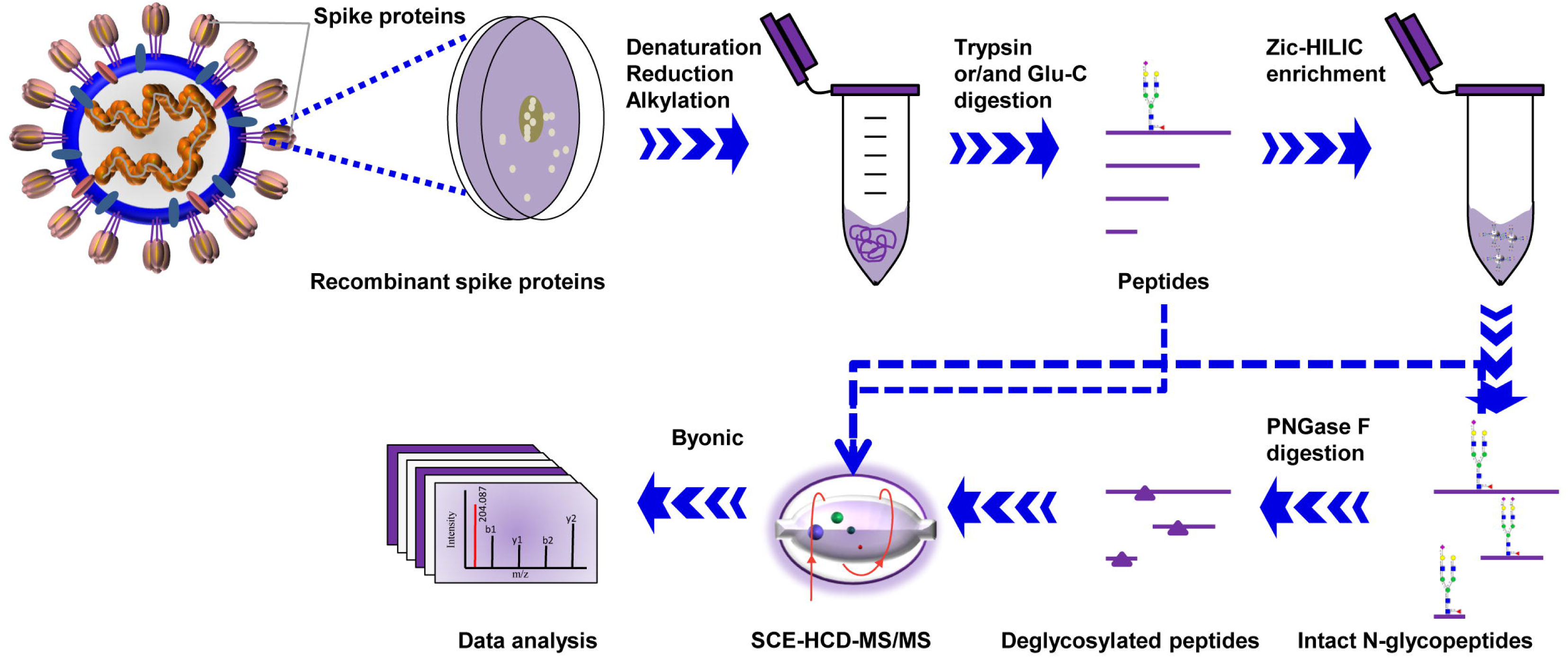
Workflow for site-specific N-glycosylation characterization of recombinant SARS-CoV-2 S proteins using two complementary proteases for digestion and an integrated N-glycoproteomic analysis.

### N-glycosite landscape of recombinant SARS-CoV-2 S proteins

The S protein contains 22 potential N-glycosites. Using our integrated analysis method, 21 glycosites were assigned unambiguously with high-quality spectral evidence (Fig. 2A and Table S1). One glycosite, N1134, was ambiguously assigned with relatively lower spectral scores (score < 200) (Fig. S2). Nevertheless, the N1134 glycosite has been observed in the Cryo-EM structure of the SARS-CoV-2 S protein(4). The relatively low spectral evidence of this glycosite indicates that a low-frequency glycosylation may occur, because our integrated methods, including glycopeptide enrichment and deglycosylation, failed to improve the spectra. Apart from the canonical N-glycosylation sequons, three non-canonical motifs of N-glycosites (N164, N334, and N536) involving N-X-C sequons were not glycosylated. Before enrichment, an average of 15 N-glycosites from trypsin-digested peptides and 13 N-glycosites from Glu-C-digested peptides were assigned. In contrast, hydrophilic enrichment resulted in a significant increase of these glycosites to 18 and 16, respectively (Table S1). To further assess the necessity for enrichment, we compared the representative spectra of one intact N-glycopeptide (N149) before and after enrichment. Without interference from the non-glycosylated peptides, the intact N-glycopeptide had more fragmented ions assigned to N-glycosites after enrichment (Fig. S3). Complementary digestion with trypsin and Glu-C promoted the confident identification of four N-glycosites (N603, N616, N709 and N717) on two intact N-glycopeptides (Table S1 and Fig. S1C). The introduction of Glu-C digestion resulted in the production of two short intact N-glycopeptides containing 23 and 36 amino acids, respectively. These peptides are more suitable for achieving better ionization and fragmentation than the long peptide of 48 and 57 amino acids obtained from trypsin digestion (Fig. S4). Deglycopeptides are suitable for verifying glycosylation sites (Fig. S5). Unexpectedly, deglycopeptide peptides led to the loss of a few glycosites, presumably because of peptide loss during deglycosylation procedures. However, almost all glycosites were confidently confirmed using trypsin and Glu-C dual digestion (Table S1).

**Figure 2.**
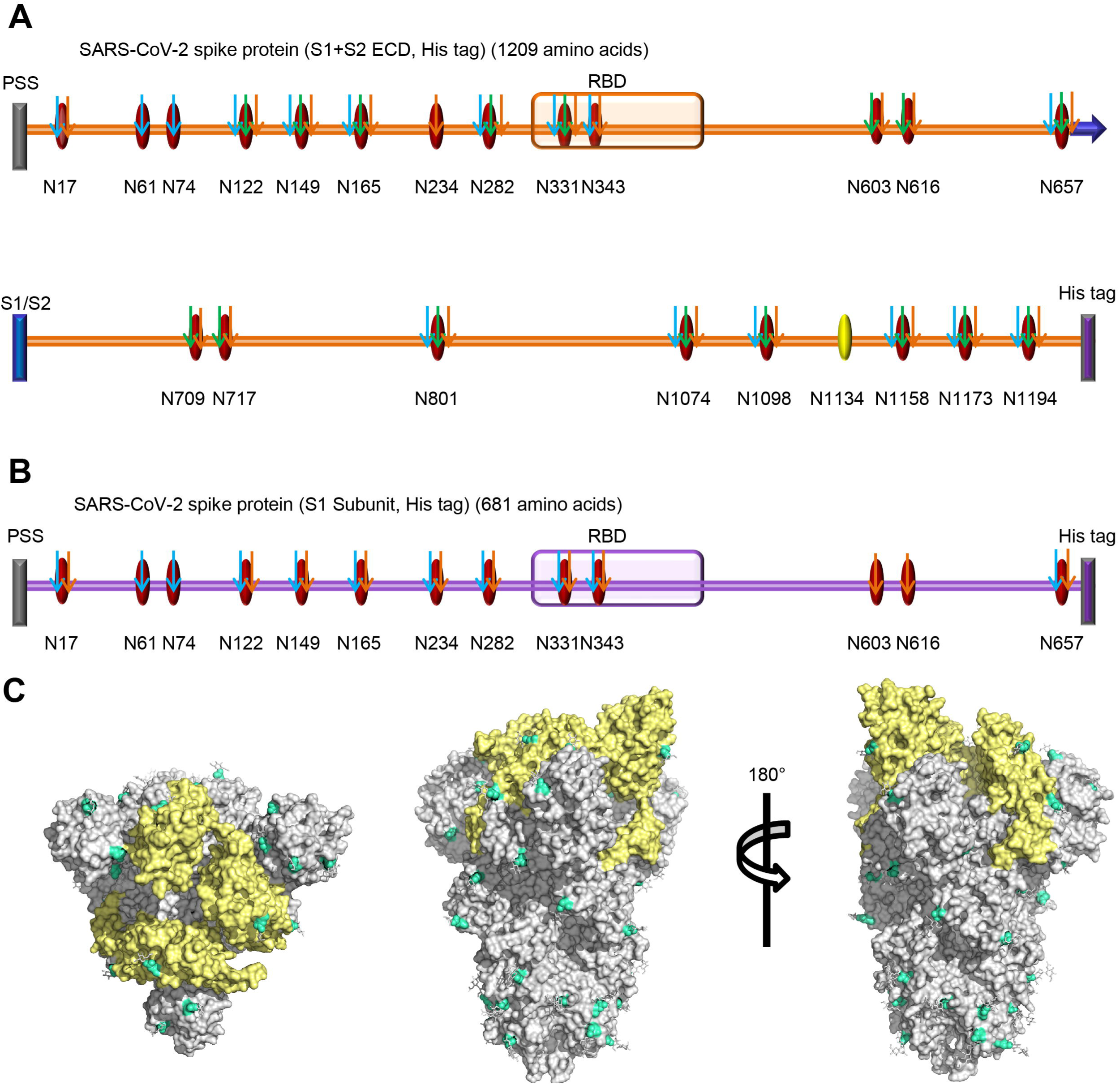
N-glycosites characterization of SARS-CoV-2 S proteins. (A and B) N-glycosites of the recombinant SARS-CoV-2 S protein or subunits expressed in insect cells (A) and human cells (B). PSS: putative signal sequence; RBD: receptor-binding domain; S1/S2: S1/S2 protease cleavage site; Oval: potential N-glycosite; Yellow oval: ambiguously assigned N-glycosite; Red oval: unambiguously assigned N-glycosite; Blue arrow: unambiguously assigned N-glycosite using trypsin digestion; Green arrow: unambiguously assigned N-glycosite using Glu-C digestion; Yellow arrow: unambiguously assigned N-glycosite using the combination of trypsin and Glu-C digestion. The unambiguously glycosite was determined by at least twice identification within each digestion list in Table S1 and Table S2. (C) N-glycosites were demonstrated in the three-dimensional structure of the SARS-CoV-2 S protein trimers (PDB code: 6VSB). RBDs, yellow; N-glycosites, blue.

For the recombinant protein S1 subunit expressed in human cells, all 13 N-glycosites were assigned unambiguously (Table S2). Finally, we profiled all 22 potential N-glycosites of S protein (Table S3 and S4). These sites were preferentially distributed in the S1 subunit of the N-terminus and the S2 subunit of the C-terminus, including two sites in the RBD (Fig. 2A and 2B). To visualize the N-glycosylation on the protein structure, all of the experimentally determined N-glycosites were hand-marked on the surface of the trimeric S protein following refinement of the recently reported SARS-CoV-2 S protein Cryo-EM structure (PDB: 6VSB) (Fig. 2C)(13).

Based on these findings, we further analyzed the conservation of glycosites among 753 SARS-CoV-2 genome sequences from the Global Initiative on Sharing All Influenza Data (GISAID) database. After the removal of redundant sequences of the S protein at the amino acid residue level, we refined 145 S protein variants. A very low frequency of alterations in 38 residue sites was found, uniformly spanning the full length of the S protein among all S variants, except for the variant D614G, which was identified at a high frequency in 98 variants (Table S5). However, nearly all of the 22 N-glycosylated sequons were conserved in the S protein, except for the loss of the N717 glycosite due to the T719A substitution in only one S variant. Following further comparison with the closely related SARS-CoV S protein(5, 29), 18 of the 22 N-glycosites were identified as conserved in the SARS-CoV-2 S protein, indicating the importance of glycosylation of the virus. Four newly arisen N-glycosites (N17, N74, N149, and N657) are located in the SARS-CoV-2 S protein S1 subunit away from the RBD. Moreover, four previously confirmed N-glycosites (N29, N73, N109, and N357) in the SARS-CoV S protein were missing in SARS-CoV-2 S, one of which (N357) lies in the RBD (Fig. S6). Additionally, two N-glycosites (N1158 and N1173) identified in SARS-CoV-2 S in this study were not detected in SARS-CoV S (N1140 and N1155) in previous studies(5, 29). Our results suggest that the preferential change of the glycosylation landscape of the S1 subunit tends to change the distribution of glycan shield, especially in the N terminal half of S1 (Fig. S6).

### Site-specific N-glycan occupancy of recombinant SARS-CoV-2 S proteins

Intact N-glycopeptide analysis can provide N-glycoproteomic information, including the composition and number of N-glycans decorating a specific N-glycopeptide or N-glycosite. The potential N-glycopeptides in the S protein sequence are shown in Fig. S1. A comparison of the intact N-glycopeptides’ spectra to the total spectra showed that the average enrichment efficiency of the Zic-HILIC materials reached up to 97%. Ultimately, hundreds of non-redundant intact N-glycopeptides were identified from the recombinant S ectodomain (Table S3) and S1 subunit (Table S4). Representative and high-quality spectra of intact N-glycopeptides are shown in Fig. S7. Following glycopeptide enrichment, the number of intact N-glycopeptides and N-glycans significantly increased (*P*<0.05) (Fig. 3A and Fig. 3B).

**Figure 3.**
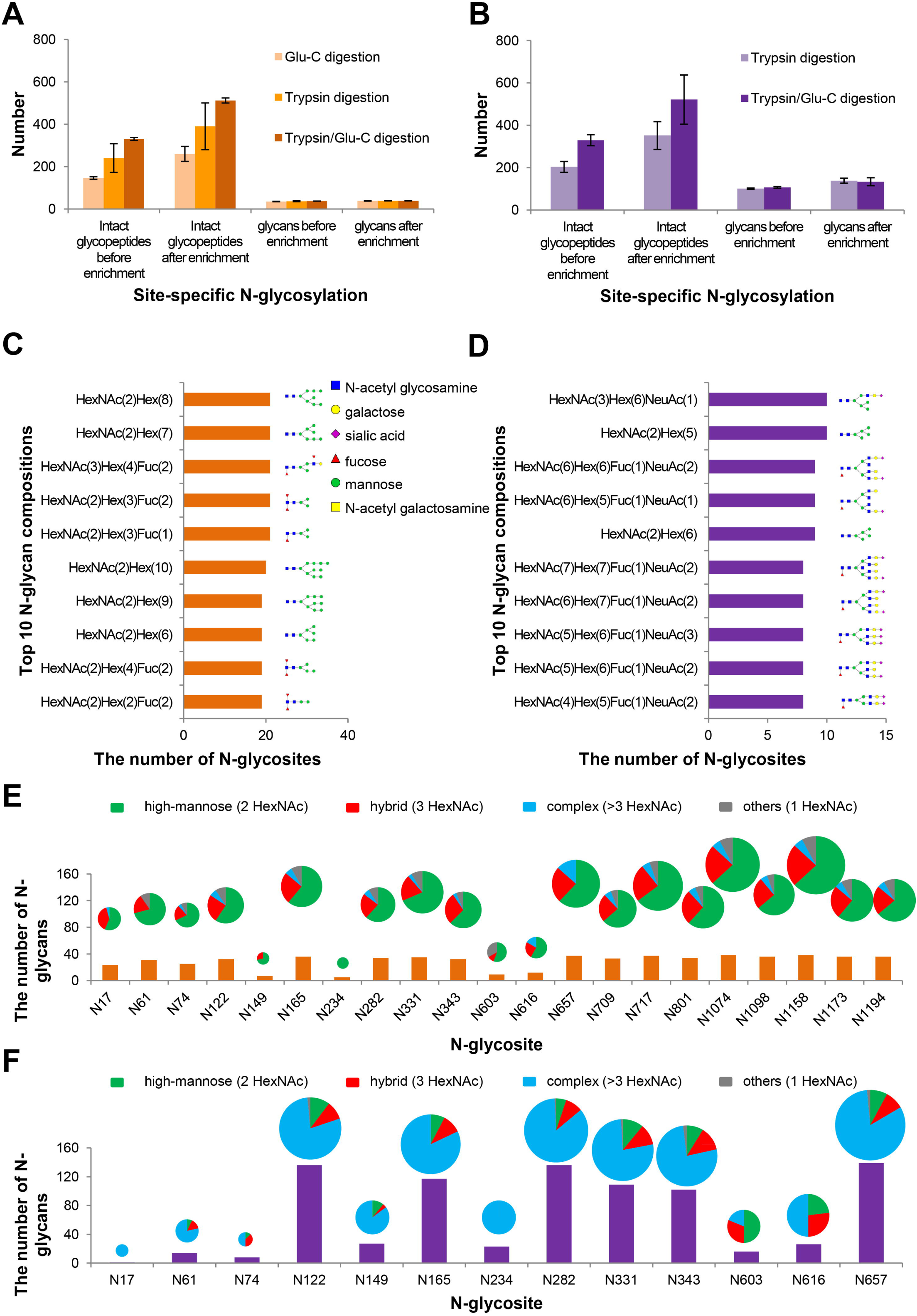
Site-specific N-glycosylation of recombinant SARS-CoV-2 S proteins. (A and B) The number of intact N-glycopeptides and N-glycans in recombinant SARS-CoV-2 S proteins expressed in insect cells (A) or human cells (B). (C and D) The numbers of the N-glycosites containing one representative N-glycan and its deduced structure from the recombinant SARS-CoV-2 S protein or subunit expressed in insect cells (C) and human cells (D). (E and F) Different types and numbers of N-glycan compositions on each N-glycosite of the recombinant SARS-CoV-2 S protein or subunit expressed in insect cells (E) or human cells (F).

Regarding the N-glycan composition, S protein expressed in insect cells had smaller and fewer complex N-glycans attached to intact N-glycopeptides than S1 subunit produced in human cells. Both recombinant products contained the common N-acetylglucosamine (HexNAc) as a canonical N-glycan characteristic (Fig. 3C and 3D). S protein expressed in insect cells were decorated with 38 N-glycans, with the majority preferentially containing oligomannose (Hex) and fucose (Fuc) (Fig. 3C and Table S3). By contrast, the S1 subunit expressed in human cells were attached to up to 157 N-glycans, mainly containing extra N-acetylglucosamine (HexNAc) and galactose (Hex), variably terminating with sialic acid (NeuAc) (Fig. 3D and Table S4). Returning to the glycosite level, most of the N-glycosites in the S protein were modified with 17–35 types of N-glycans and classified into a high proportion of high-mannose N-glycans (∼65%) and a lower proportion (∼23%) of hybrid N-glycans. Almost all N-glycosites contained no more than 10% of complex N-glycans (Fig. 3E). For the S1 subunit expressed in human cells, the occupancy of N-glycans on each N-glycosite was quite nonuniform. Surprisingly, six N-glycosites (N122, N165, N282, N331, N343, and N657) were decorated with markedly heterogeneous N-glycans of up to 139 types. The average occupancies of all glycosites presented as an overwhelming proportion (∼75%) of complex N-glycans and a small proportion of hybrid (∼13%) or high-mannose (∼12%) N-glycans (Fig. 3F). The glycan occupancy on two N-glycosites (N331 and N343) of RBD were identified (Fig. 3E and 3F). The high occupancy of RBD glycosites by various N-glycan compositions implies that N-glycosylation might be associated with the recognition of RBD to ACE2 receptor, since the interaction between RBD and ACE2 mainly depends on polar residue interactions(14). Our results suggest that S proteins expressed in different cells display distinct N-glycosylation patterns. In particular, the glycosylation of the S protein in human cells exhibits remarkable heterogeneity on N-glycosites. However, the N-glycan types on each glycosite is primarily determined by the host cells rather than the location of different glycosites (Fig. 3E-3F, Fig. S8).

### Site-specific N-glycan occupancy of recombinant SARS-CoV-2 S protein RBD

To confirm site-specific N-glycan occupancy and exploit the potential impact on N-glycosylation by different protein sizes, recombinant RBDs (residues 319–541) from both human and insect cells were further analyzed (Table S6). The representative glycan compositions and deduced structures are shown on each site (Fig. 4A). Intriguingly, the number of glycan compositions and their types on each glycosite (Fig. 4B and 4C) are very close to those found in the S ectodomain and S1 subunit (Fig. 3E and Fig. 3F). The human cell-produced RBDs displayed more N-glycan compositions and complex glycan types, compared to insect cell-expressed proteins (Fig. 4B and 4C). Moreover, more than 80% of glycan compositions are identical at each site among different lengths of insect cell-expressed proteins (Fig. 4D). Similarly, over 75% of the glycan compositions at each site were found to be shared by human cell-produced products (Fig. 4E). The N-glycosylation of RBDs was verified by SDS-PAGE (Fig.S9). These results suggest that the N-glycan compositions are conserved among different sizes of RBD proteins. Taken together, our data reveal the regular heterogeneity of N-glycan compositions at each site of the S protein subunits, primarily depending on host cells and glycosites. Intriguingly, the N-glycan types on S protein subunits are predominantly determined by host cells, regardless of the location of glycosites.

**Figure 4.**
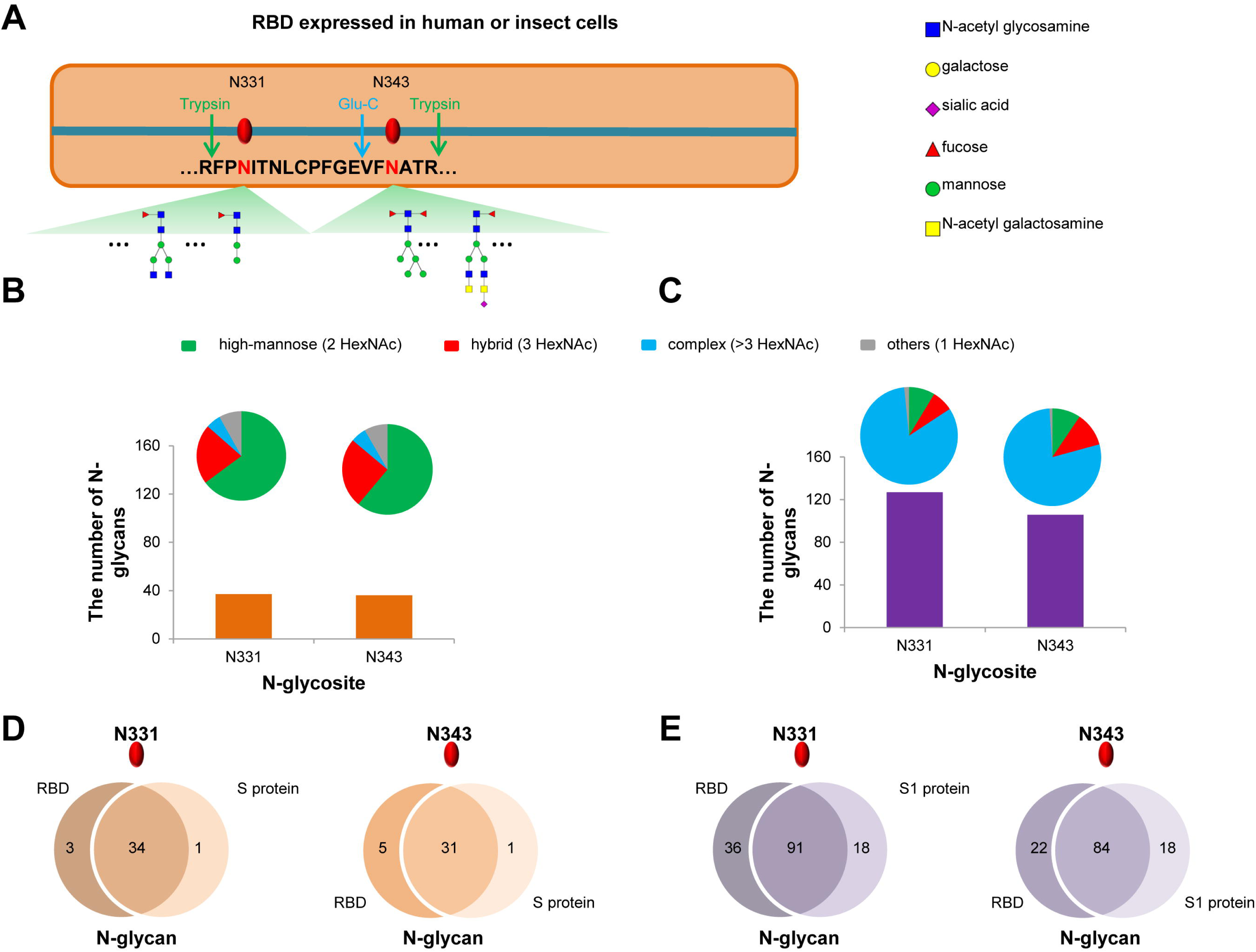
Site-specific N-glycosylation profile of RBDs expressed in human and insect cells. (A) The glycosites and deduced representative N-glycans on N331 and N343 of RBD. (B and C) Different types and numbers of N-glycans on N331 and N343 of RBD expressed in insect cells (B) or human cells (C). (D) Comparison of N-glycans on N331 and N343 between RBD and S ectodomain expressed in insect cells. (E) Comparison of N-glycans on N331 and N343 between RBD and S1 subunit expressed in human cells.

## DISCUSSION

The global outbreak and rapid spread of COVID-19 caused by SARS-CoV-2 urgently call for specific prevention and intervention measures(30). The development of preventative vaccines and neutralizing antibodies remains a chief goal in the efforts to control viral spread and stockpile candidates for future use. However, this work greatly relies on the understanding of the antigen structure and state of glycosylation for the rational determination of accessible epitopes. The S protein is posited to be the main or even the only antigen on viral surfaces for priming the immune system to produce an effective response (21, 26). Previous studies have revealed the structural information of the SARS-CoV-2 S protein and found the coverage of N-glycans (4, 13). In this study, we profiled the site-specific N-glycosylation of the recombinant SARS-CoV-2 S protein. All potential N-glycosites on the S protein were identified experimentally. The N-glycan compositions and types on S protein subunits were revealed among different host cells. Our data provide a large-scale N-glycosylation information of the S protein and present a promising prospect for developing a vaccine and therapeutic antibodies.

N-glycosylation is a common feature of the viral envelope proteins, including those of HIV-1, Lassa virus, hepatitis C virus, Epstein–Barr virus and influenza A virus. Glycosylation promotes proper glycoprotein folding; however, the glycans obstruct receptor binding and proteolytic processing during antigen presentation(21, 23, 31). Characterization of the landscape of N-glycans on the SARS-CoV-2 S protein is crucial for promoting immunogen design and prevention of the potential viral evasion of intervention measures(5, 21, 32). Precise characterization of intact N-glycopeptides can reveal the occupancy of each glycosite by different glycoforms(33, 34). In this study, all 22 N-glycosites of the S protein were identified (Fig. 2A and 2B). By comparison, the alteration of N-glycosites among SARS-CoV-2 and SARS-CoV S proteins focused on glycosites in the S1 subunit (Fig. S6). The N-glycosites located in the S2 subunit are completely conserved, and seven out of nine sites, along with glycans have been disclosed by previous studies(4, 5). Moreover, we found that N-glycosites in the S protein were highly preserved among 145 SARS-CoV-2 S protein variants (Table S5), which is advantageous for circumventing potential viral immune evasion from the vaccines and neutralizing antibodies currently being developed.

Glycosylation of proteins is intricately processed by various enzymes coordinated in the endoplasmic reticulum and Golgi apparatus. The glycan composition and structure decoration at specific sites occurred in a non-templated manner governed by host cells, thus frequently resulting in a heterogeneous glycan occupancy on each glycosite(35). Glycosylation processing in insect and human cells can yield a common intermediate N-glycan, which will further elongate in human cells but is only trimmed in insect cells to form end products(27). Consequently, the S protein expressed in human cells displayed a larger size and a much higher proportion of complex N-glycans than that expressed in insect cells, owing to the additional elongation of the glycan backbone with multiple oligosaccharides (Fig. 3C-3F). Our findings are in line with those of previous studies, that demonstrated the predominant complex of N-glycans attached to MERS-CoV and SARS-CoV proteins (4, 5, 36). Moreover, two recent studies have revealed that the complex N-glycans dominate the glycosites on human cell-expressed SARS-CoV-2 S protein(37, 38). In contrast, S protein expression in insect cells led to a high ratio of high-mannose N-glycans (Fig. 3C and 3E), which has also been found in the insect cell-produced HCoV-NL63 S protein, despite expression in a different insect cell, the *Drosophila S2* cell, in the previous study(21).

Notably, a low ratio (∼12%) of high-mannose glycans ubiquitously exist on human cell-expressed S protein (Fig. 3F). The HIV envelope glycoprotein gp120 is heavily decorated with the immature intermediate, high-mannose glycans. The high-density glycans surrounding HIV glycoproteins limit the accessibility of glycan biosynthetic processing enzymes, terminating the synthesis of more complex end products(23, 32). By contrast, the high ratio of complex N-glycans in the SARS-CoV-2 S protein were successfully processed on most glycosites by the enzymes, without extensive obstruction by the on-going synthesis of glycan shields (Fig. 3F and Fig. 4C). Therefore, we posit that the glycan coverage on the SARS-CoV-2 S protein could leave relatively accessible antigens and epitopes, although the complex N-glycans might mask some surface immunogens. These features may provide a promising landscape on the SARS-CoV-2 S protein for immune recognition. This potential is bolstered by the findings that the convalescent sera from COVID-19 patients contains antibodies against the SARS-CoV-2 S protein(7, 10). A previous study on SARS-CoV has revealed that the oligomannose on the S protein can be recognized by mannose-binding lectin (MBL) and may interfere with viral entry into host cells by inhibition of S protein function(39). Besides the direct neutralization effect, MBL, as a serum complement protein, can initiate the complement cascade. Complement hyper activation in lung tissues of COVID-19 patients has been revealed by a recent preprint study(40). However, it remains unknown whether MBL can bind to oligomannose on SARS-CoV-2 S protein to impact the viral spread or initiate complement activation in patients. Glycans also play crucial and multifaceted roles in B cell and T cell differentiation via cell-surface or secreted proteins, including selectins, galectins, and siglecs, which can further connect SARS-CoV-2 to immune response and immune regulation(35). In particular, the complex N-glycans are ligands for galectins, which are able to engage different glycoproteins to regulate immune cell infiltration and activation upon virus infection(41, 42). These mechanisms underlying SARS-CoV-2 infection and spread are worth further clarification based on a detailed analysis of clinical characteristics in humoral and cellular immunity.

The remarkable heterogeneity of N-glycosylation in the S protein subunit expressed in human cells was revealed in our study (Fig. 3F). By contrast, the N-glycosylation of the S protein subunit in insect cells showed less heterogeneity and complexity than that of human cell-derived proteins (Fig. 3E). Moreover, the site-specific glycan occupancy tended to be identical in the same host cell, regardless of protein length (Fig. 4D and 4E). These results indicate the N-glycan compositions and types on S protein largely attribute to different host cells with the differential processing pathways of glycosylation. We can expect that the native N-glycosylation profile of the SARS-CoV-2 S protein in humans tends to be consistent with that of the recombinant protein expressed in human cells, unless the virus buds off early in the glycosylation processing pathway and produces immature glycans(27, 35, 43).

Intriguingly, the immature N-glycans such as high-mannose are regarded as “non-self” glycans (43, 44). Therefore, the insect cell-expressed recombinant antigens decorated with paucimannose and high mannose are more immunogenic in mice than those produced in human cells(45, 46). In contrast, the vaccine antigens produced from mammalian cells do not always induce a strong humoral immune response in mice, because of the complex-type N-glycans(47). To prime strong humoral immunity upon vaccination against SARS-CoV-2, insect cell-produced antigens with less complex N-glycans could be one of the candidates for the development of vaccines and neutralizing antibodies. Apart from amino acid epitopes, the glycopeptide can be presented by major histocompatibility complex (MHC) and recognized by a CD4+ T-cell population to help B cells produce antibodies against glycans. The glycoconjugate has been used to boost the immune response against infections(31, 32). The insect cell-produced S protein subunits could prime protective immunity against the “non-self” oligomannose N-glycans, in case of the immature N-glycans linked to the native envelope proteins of SARS-CoV-2, which seems to occur in the SARS-CoV replication(43). In contrast, the human cell-expressed S protein subunits as vaccines mimic the “self” glycans in human, which are not expected to boost immune response to the glycoantigens. However, the remaining accessible and non-glycosylated regions can serve as the antigens and epitopes. The rational design of antigens to prime potent and broad immune responses against accessible epitopes on SARS-CoV S protein is essential and promising.

The RBD-containing subunit is an ideal immunogen since antibodies against the receptor-binding motif within RBD could directly block the engagement of S protein to the receptor and inhibit viral infections of host cells. Vaccination with SARS-CoV RBD has been demonstrated to induce potent and long-term immunity in animal models(48). Meanwhile, the subunit vaccines are posited to minimize the potentially undesired immunopotentiation of the full-length S protein, which might induce severe acute injury in the lungs(49). Intriguingly, SARS-CoV-2 is missing one N-glycosite in RBD compared to SARS-CoV. The remaining two N-glycosites were outside of the motifs essential for direct interaction with the ACE2 receptor(14) (Fig. 2C). The glycan compositions of RBD are highly identical in the same host cell, regardless of the length of the RBD-containing proteins (Fig. 4B-4E). These features of the RBD, along with its highly exposed structure, endow more antigens and accessible epitopes for vaccine design and immune recognition. The RBD-containing proteins, especially the insect cell-expressed products, could become promising candidates for SARS-CoV-2 vaccine development. However, drug discovery related to glycosylation inhibition is supposed to be performed based on human cell-expressed products.

In this study, we decoded a global and site-specific profile of N-glycosylation on SARS-CoV-2 S proteins expressed from insect and human cells, revealing a regular heterogeneity in N-glycan composition and site occupancy. All glycosites were conserved among the 753 public SARS-CoV-2 genome sequences. In conclusion, our data indicate that differential N-glycan occupancies among distinct host cells might help elucidate the infection mechanism and develop an effective vaccine and targeted drugs. Nevertheless, the implication of S protein site-specific N-glycosylation in immunogenicity, receptor binding, and viral infectivity should be investigated further.

## Abbreviations

ACE2: angiotensin-converting enzyme II
Cryo-EM: cryoelectron microscopy
E: envelope protein
HCoV-NL63: human coronavirus NL63
M: membrane protein
MS: mass spectrometry
MERS-CoV: Middle East respiratory syndrome coronavirus
N: nucleocapsid protein
RBD: receptor-binding domain
S: spike protein
SARS-CoV-2: severe acute respiratory syndrome coronavirus
SCE: stepped collision energy
Zic-HILIC: zwitterionic hydrophilic interaction liquid chromatography

## Acknowledgments

We would like to express our special thanks to the National Center for Protein Sciences Beijing, Professor Catherine E. Costello in Boston University School of Medicine, and the COVID-19 Mass Spectrometry Coalition for sharing knowledges in glycopeptide analysis. Thanks to Beijing Sino Biological Inc for providing high-quality recombinant proteins for this project.

## Data Availability

The raw MS data have been deposited to the ProteomeXchange Consortium via the PRIDE partner repository with the dataset identifier PXD018506.

## Author Contributions

H.Y., M.G., D.D., X.L., Y. C. and J.C. directed and designed research; Y.Z. and W.Z. directed and performed analyses of mass spectrometry data; Y. Z., H.Y. and S.W. adapted algorithms and software for data analysis; Y.Z. and T.S. coordinated acquisition, distribution and quality evaluation of samples; Y.Z. and H.Y. wrote the manuscript.

## Supplementary Figures and Tables

**Supplementary Figure S1**. The theoretical of intact N-glycopeptides of S protein derived from the digestion using trypsin (A) or Glu-C (B) alone or in combination (C). Red letter: N-glycosites; Green letter: trypsin cutting sites; Blue letter: Glu-C cutting sites; Underline: theoretical N-glycopeptides without missing cleavage sites.

**Supplementary Figure S2**. The spectrum of intact N-glycopeptides with the ambiguously assigned N-glycosite (N1134).

**Supplementary Figure S3**. Comparison of the spectra of the intact N-glycopeptide (N149) before (A) and after (B) enrichment.

**Supplementary Figure S4**. Comparison of the spectra of intact N-glycopeptides (N709 and N717) after trypsin digestion (A) and Glu-C digestion (B).

**Supplementary Figure S5**. Comparison of the spectra of intact N-glycopeptides (N709 and N717) (A) and deglycopeptides (B).

**Supplementary Figure S6**. Comparison of the N-glycosites on the SARS-CoV-2 and SARS-CoV spike proteins.

**Supplementary Figure S7**. Representative and high-quality spectra of intact N-glycopeptides and deglycosylated peptides.

**Supplementary Figure S8**. Microheterogeneity and macroheterogeneity of the N-linked glycopeptides of the S protein.

**Supplementary Figure S9**. SDS-PAGE analysis of RBDs expressed in insect and human cells.

**Table S1**. Glycoproteomic identification of the N-glycosites on recombinant SARS-CoV-2 S protein expressed in insect cells.

**Table S2**. Glycoproteomic identification of the N-glycosites on recombinant SARS-CoV-2 S protein expressed in human cells.

**Table S3**. Site-specific N-glycosylation characterization of recombinant SARS-CoV-2 spike protein expressed in insect cells.

**Table S4**. Site-specific N-glycosylation characterization of recombinant SARS-CoV-2 spike protein expressed in human cells.

**Table S5**. Mutation frequency of SARS-CoV-2 spike protein.

**Table S6**. Intact N-glycopeptides of the recombinant RBD of the SARS-CoV-2 spike protein.

