## Supplementary material for "Site-specific N-glycosylation Characterization of Recombinant SARS-CoV-2 Spike Proteins": Supporting_Information.docx

**Running title:** N-glycosylation of SARS-CoV-2 Spike Protein

***Corresponding Authors:** Hao Yang, PhD, Associate Researcher, Key Lab of Transplant Engineering and Immunology, West China Hospital, Sichuan University

**Address:** No. 1, Keyuan 4th Road, Gaopeng Avenue, Hi-tech Zone, Chengdu 610041, China. Phone: +86-28-85164031; Fax: +86-28-85164031;

**Supplementary Figures**

**Supplementary Figure S1.** The theoretical of intact N-glycopeptides of S protein derived from the digestion using trypsin (A) or Glu-C (B) alone or in combination (C). Red letter: N-glycosites; Green letter: trypsin cutting sites; Blue letter: Glu-C cutting sites; Underline: theoretical N-glycopeptides without missing cleavage sites.

**Supplementary Figure S2.** The spectrum of intact N-glycopeptides with the ambiguously assigned N-glycosite (N1134).

**Supplementary Figure S3.** Comparison of the spectra of intact N-glycopeptides (N149) before (A) and after (B) enrichment.

**Supplementary Figure S4.** Comparison of the spectra of intact N-glycopeptides (N709 and N717) from trypsin digestion (A) and Glu-C digestion (B).

**Supplementary Figure S5.** Comparison of the spectra of intact N-glycopeptides (N122) (A) and deglycopeptides (B).

**Supplementary Figure S6.** Comparison of the N-glycosites on SARS-CoV-2 and SARS-CoV spike proteins.

**Supplementary Figure S7.** Representative and high-quality spectra of intact N-glycopeptides and deglycosylated peptides.

**Supplementary Figure S8.** Microheterogeneity and macroheterogeneity of the N-linked glycopeptides of the S protein.

**Supplementary Figure S9.** SDS-PAGE analysis of RBDs expressed in insect and human cells.

**Supplementary Figure S1.** The theoretical of intact N-glycopeptides of S protein derived from the digestion using trypsin (A) or Glu-C (B) alone or in combination (C). Red letter: N-glycosites; Green letter: trypsin cutting sites; Blue letter: Glu-C cutting sites; Underline: theoretical N-glycopeptides without missing cleavage sites.

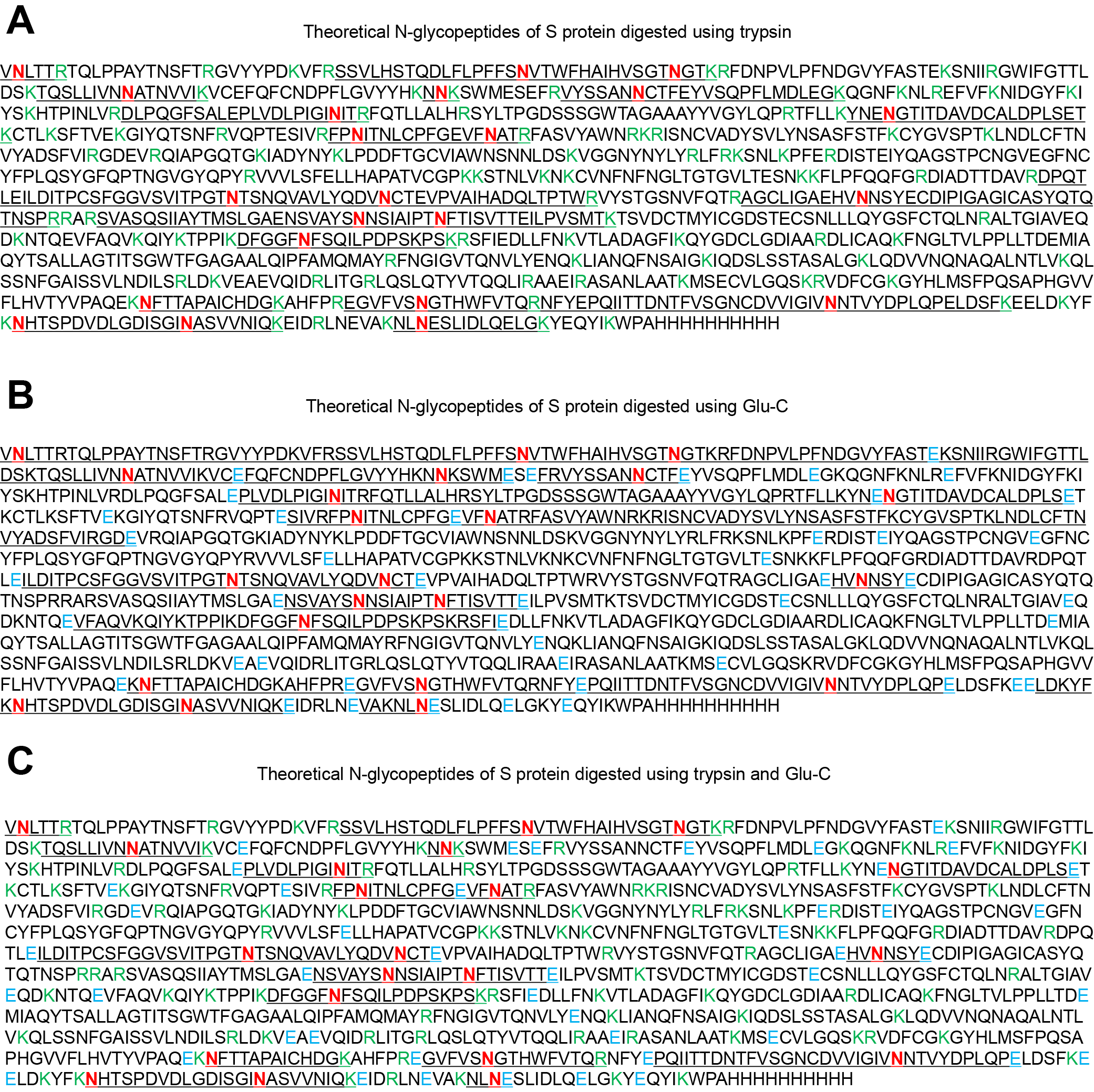

**Supplementary Figure S2.** The spectrum of intact N-glycopeptides with the ambiguously assigned N-glycosite (N1134).

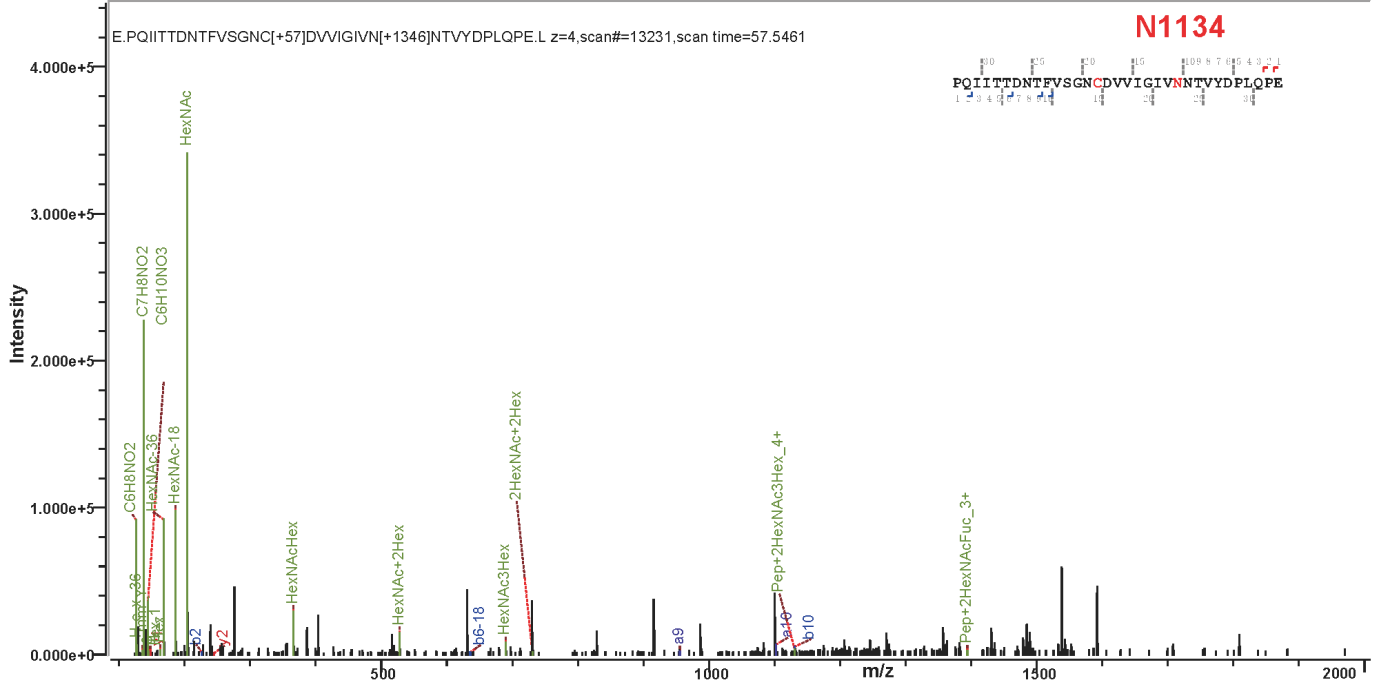

**Supplementary Figure S3.** Comparison of the spectra of the intact N-glycopeptide (N149) before (A) and after (B) enrichment.

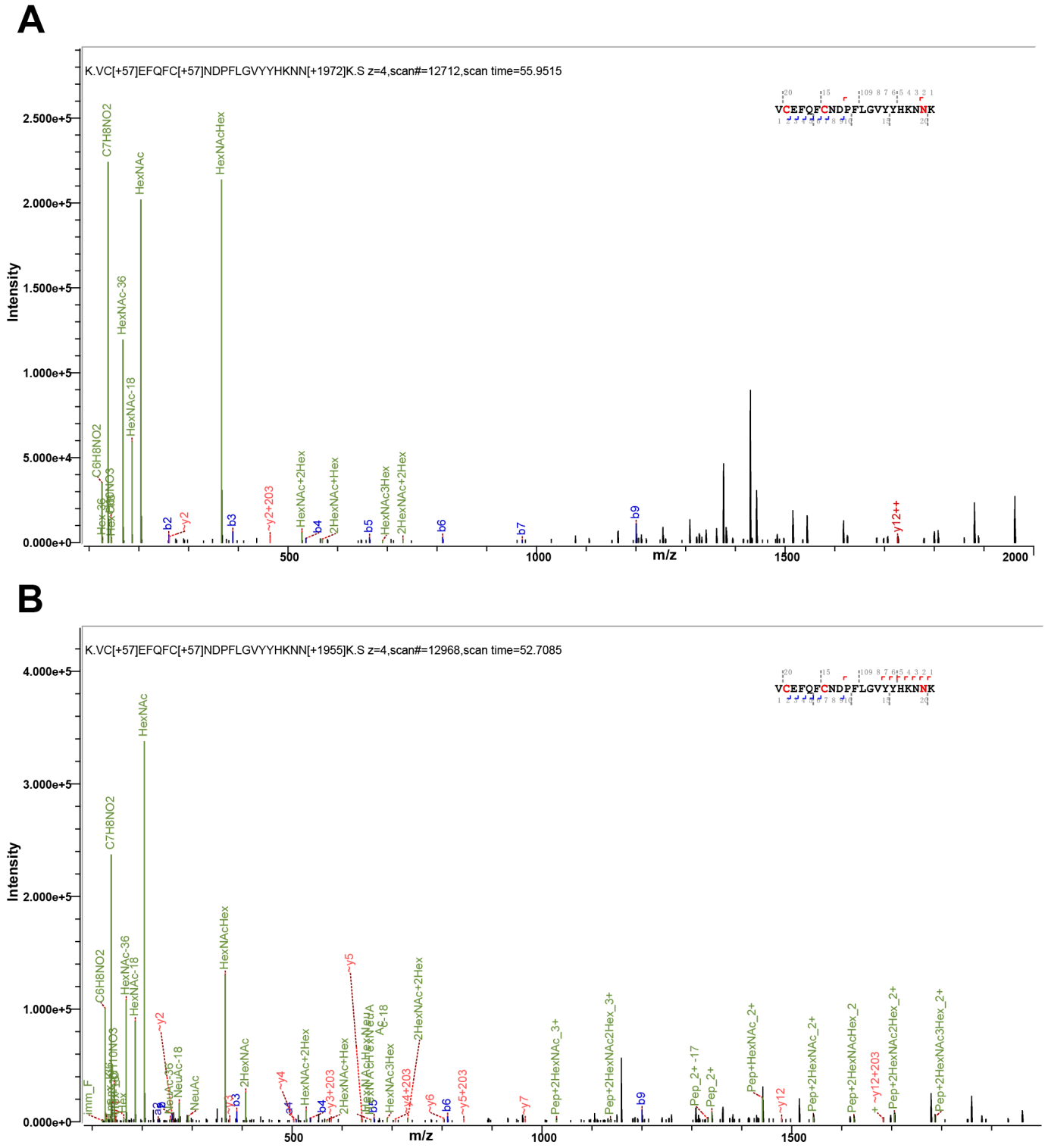

**Supplementary Figure S4.** Comparison of the spectra of intact N-glycopeptides (N709 and N717) after trypsin digestion (A) and Glu-C digestion (B).

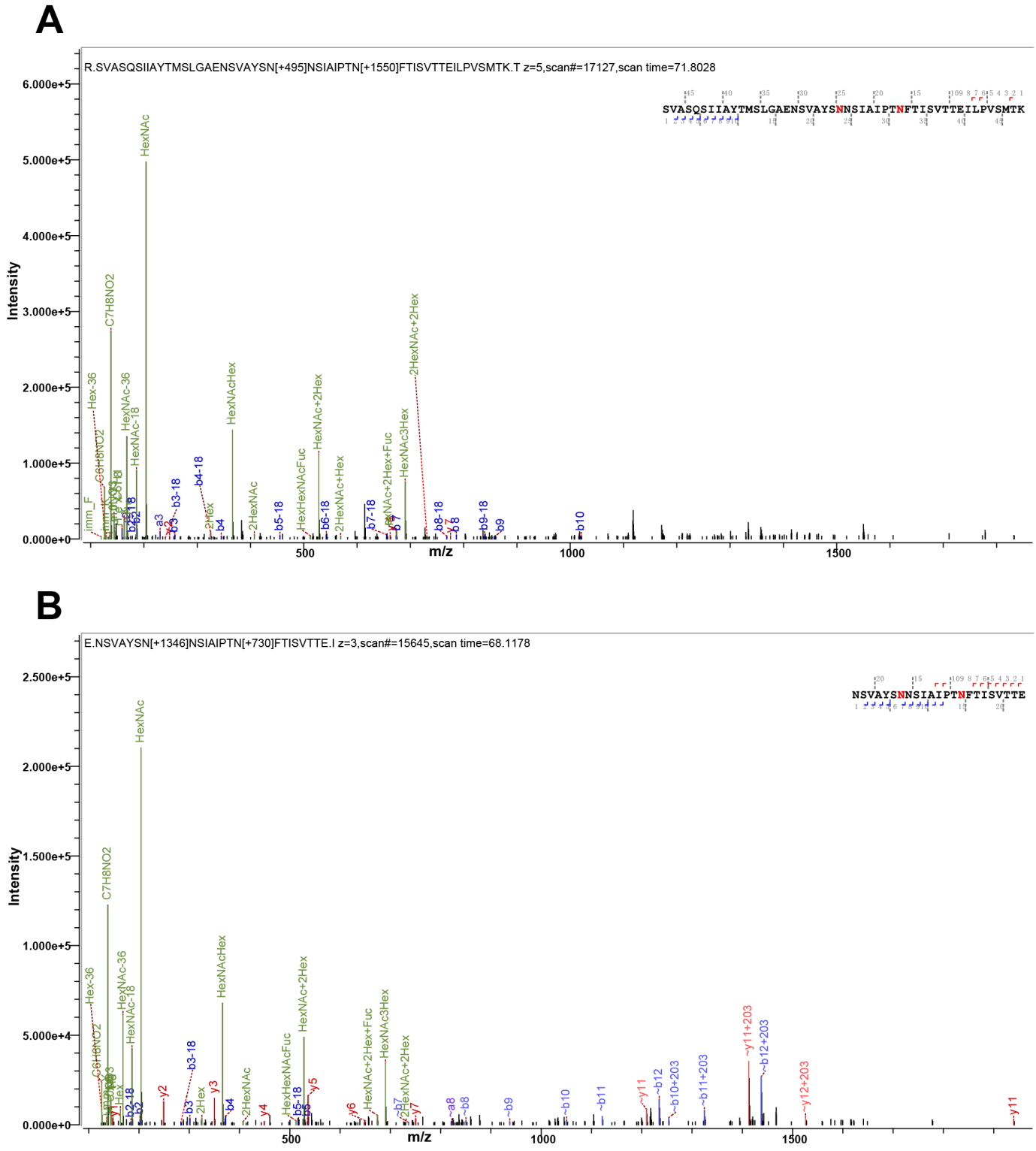

**Supplementary Figure S5.** Comparison of the spectra of intact N-glycopeptides (N709 and N717) (A) and deglycopeptides (B).

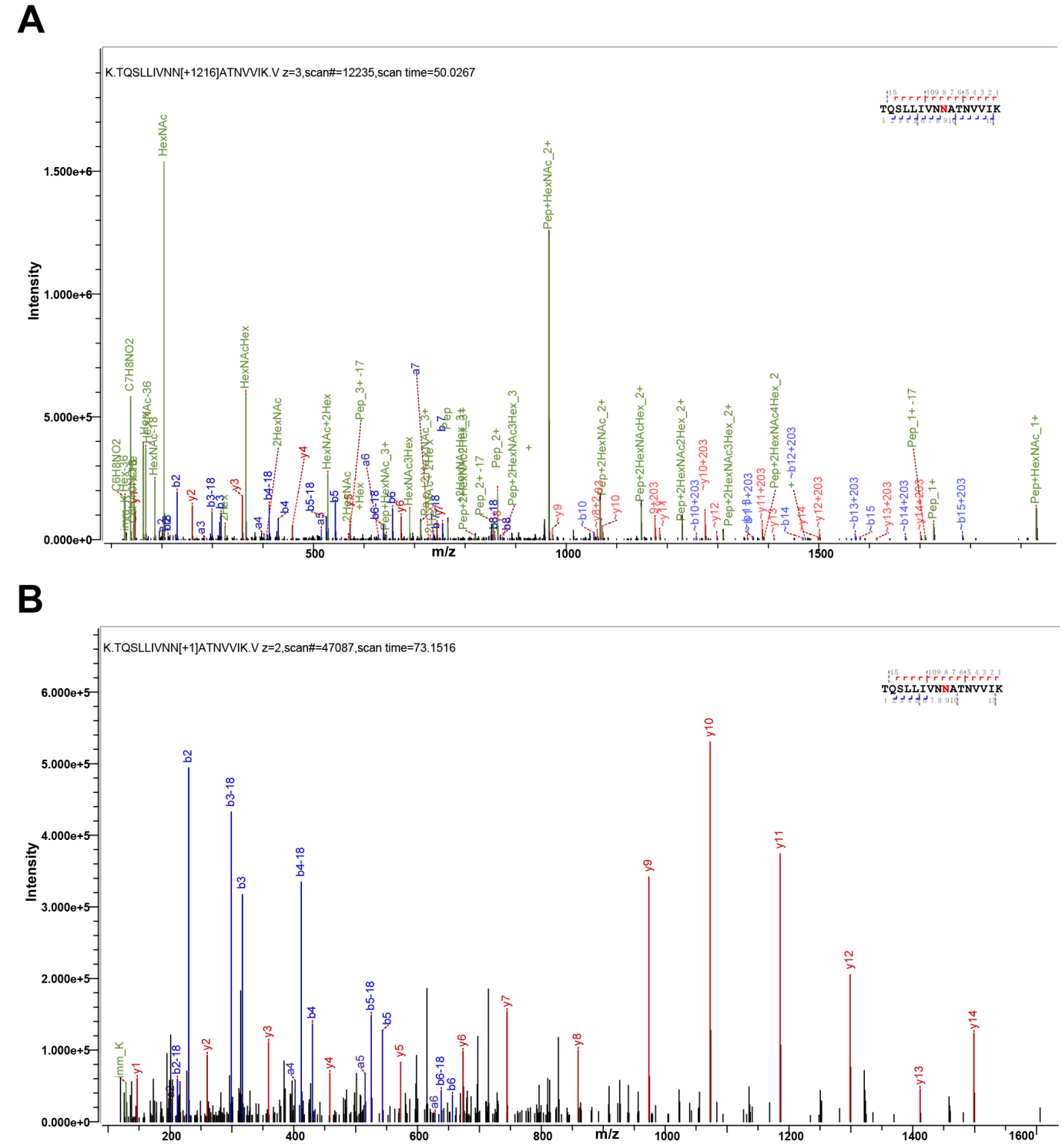

**Supplementary Figure S6.** Comparison of the N-glycosites on the SARS-CoV-2 and SARS-CoV spike proteins.

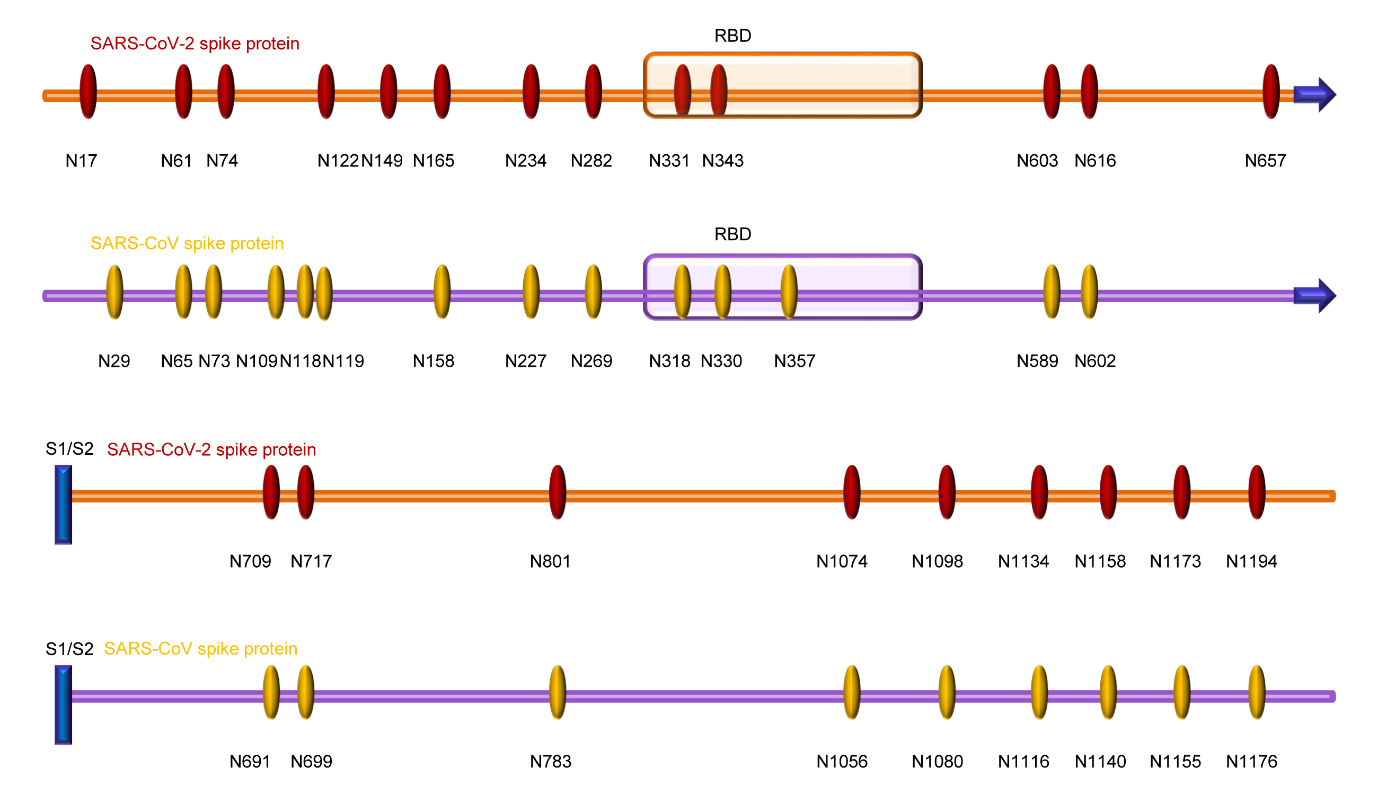

**Supplementary Figure S7.** Representative and high-quality spectra of intact N-glycopeptides and deglycosylated peptides.

**Spectra of intact N-glycopeptides from recombinant SARS-CoV-2 spike proteins expressed in insect cells**

**N17**

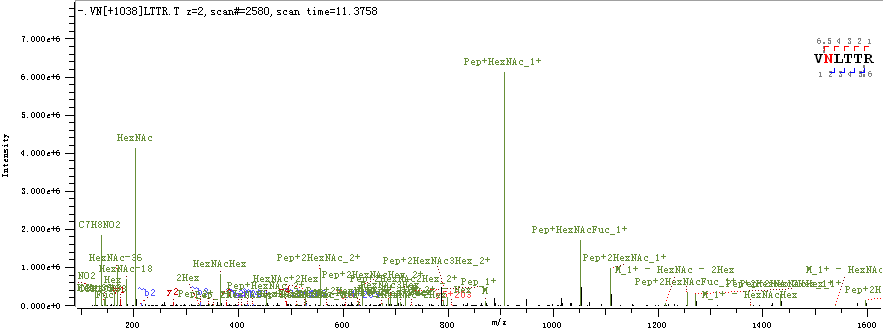

**N61 & N74**

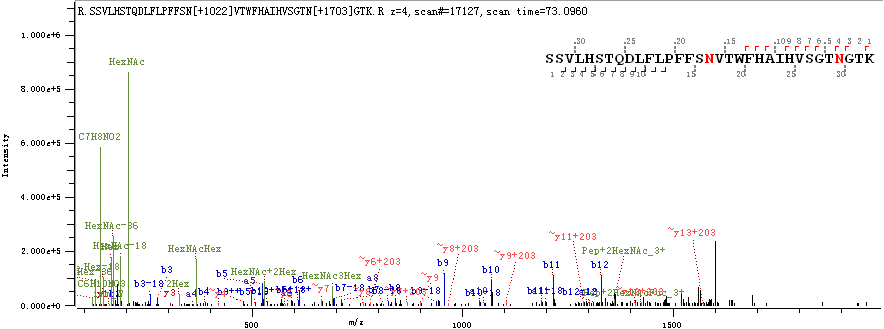

**N122**

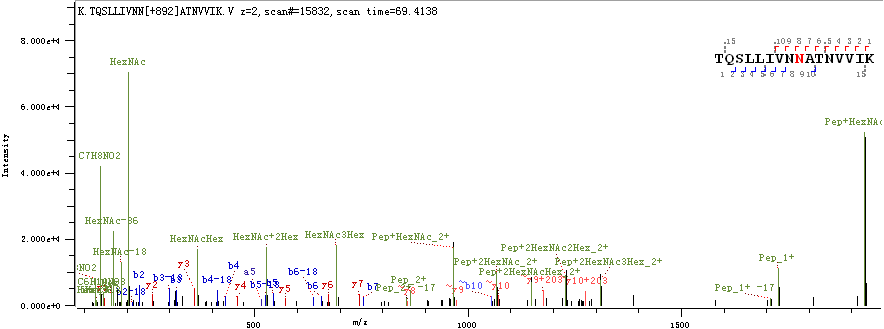

**N149**

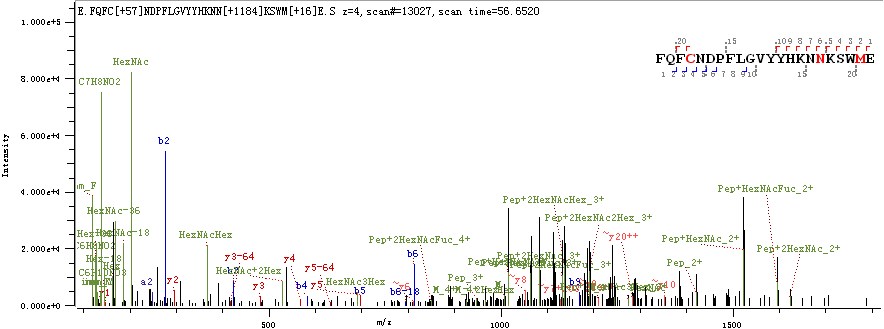

**N165**

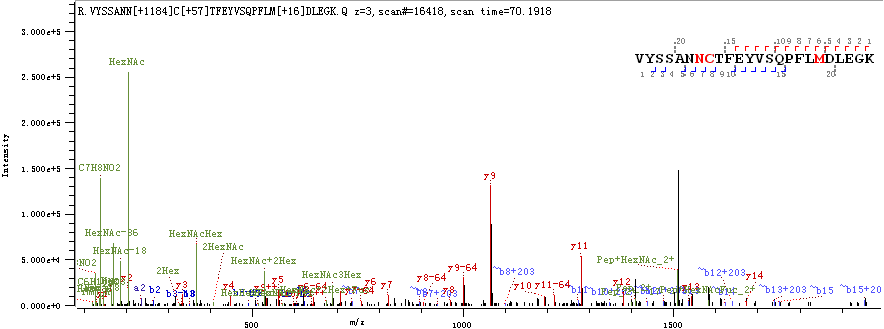

**N234**

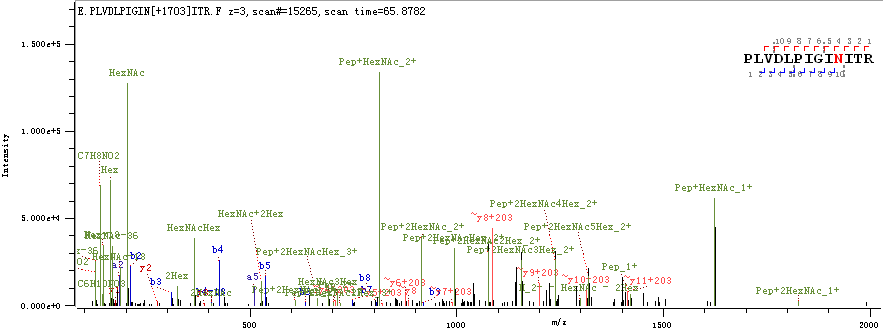

**N282**

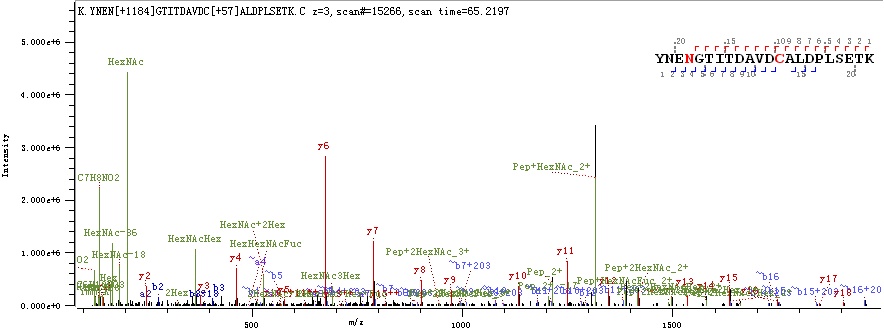

**N331 & N343**

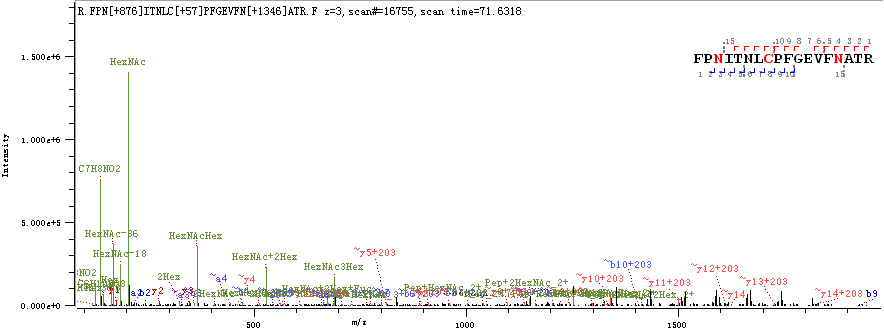

**N603 & N616**

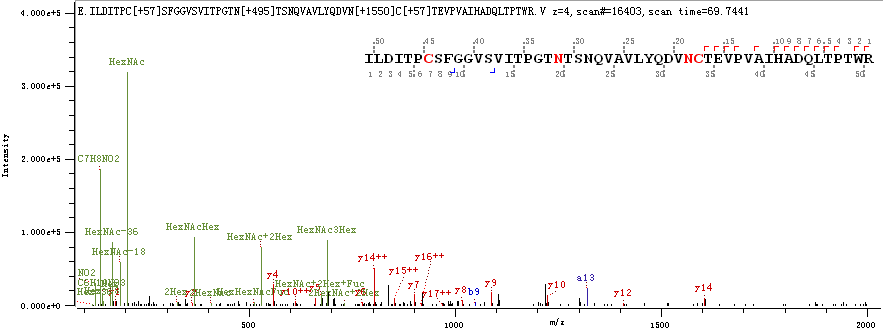

**N657**

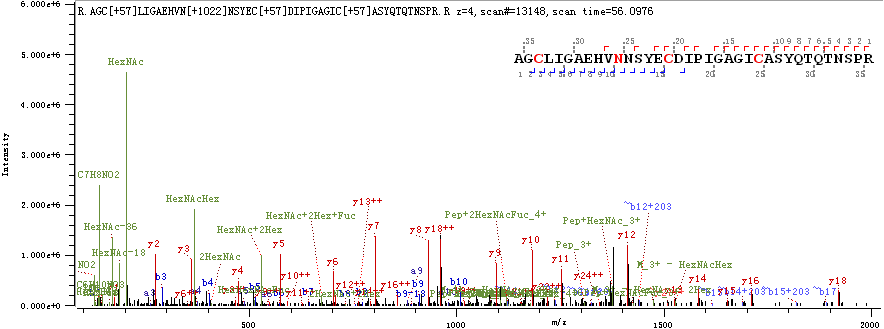

**N709 & N717**

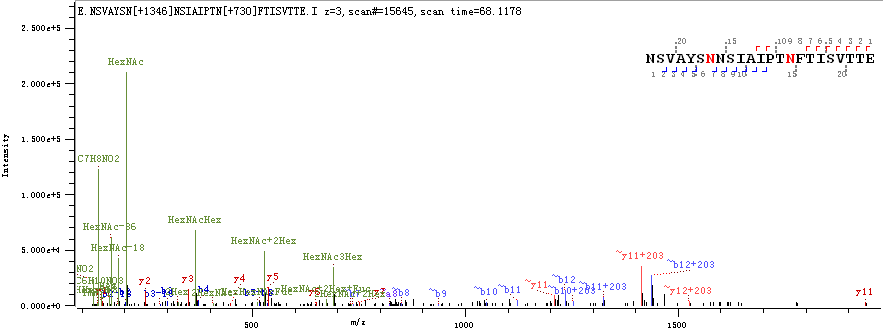

**N801**

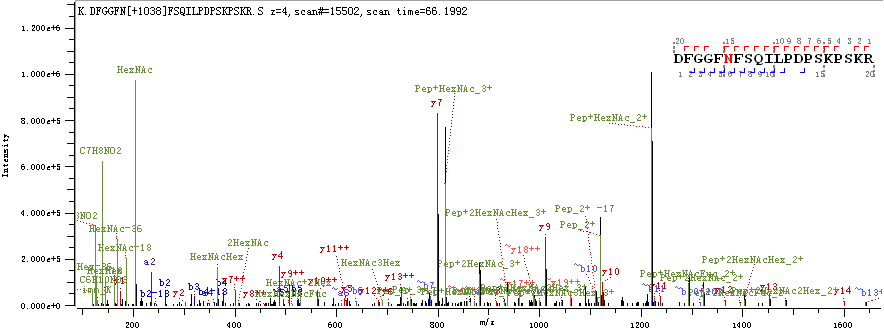

**N1074**

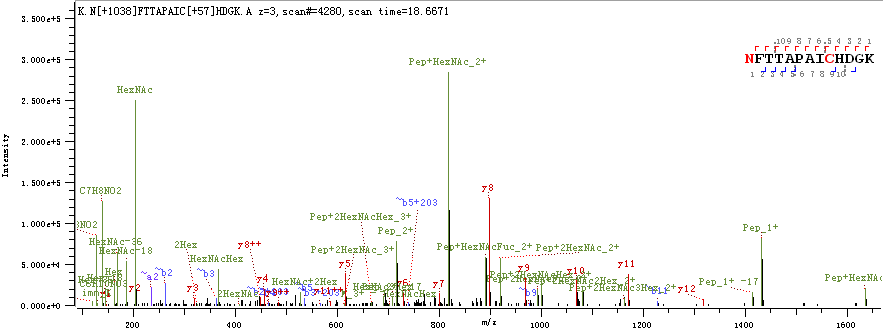

**N1098**

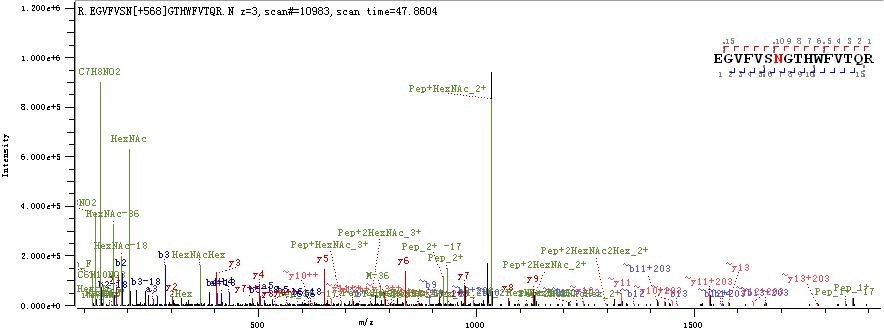

**N1158 & N1173**

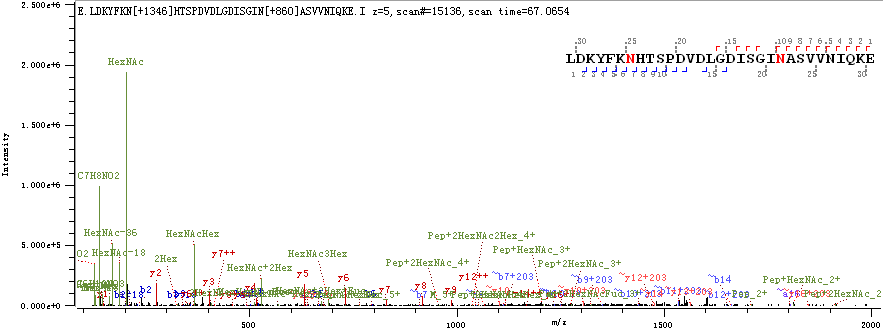

**N1194**

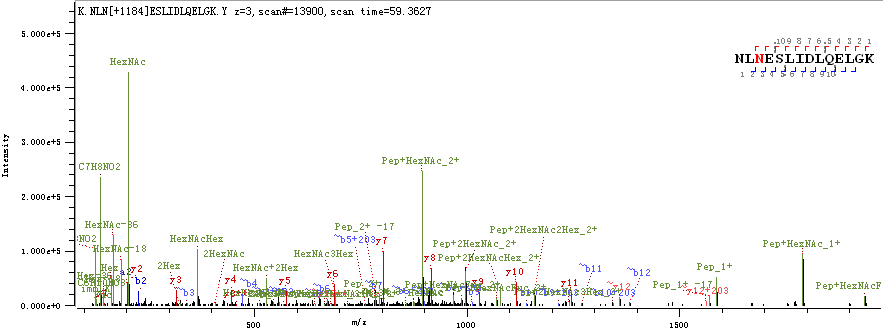

**Spectra of deglycosylated peptides from recombinant SARS-CoV-2 spike proteins expressed in insect cells**

**N17**

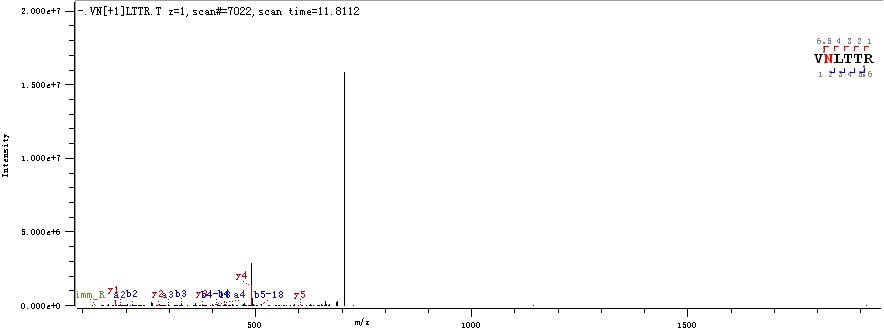

**N122**

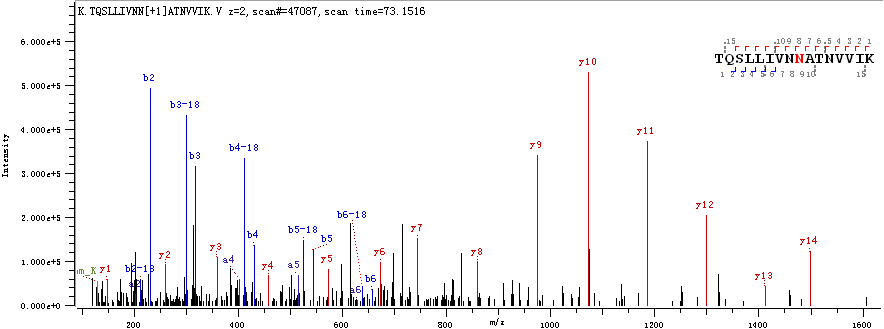

**N165**

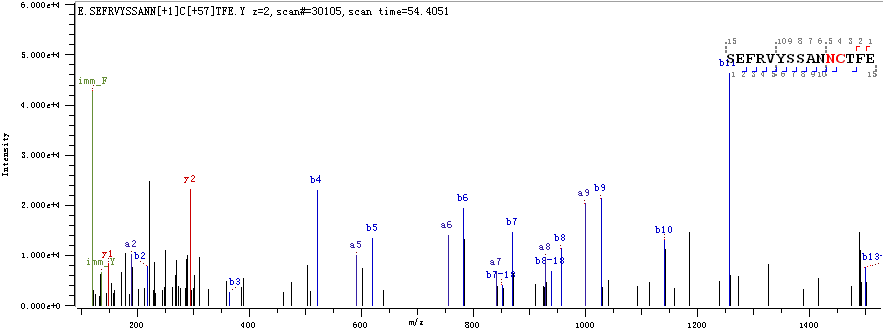

**N234**

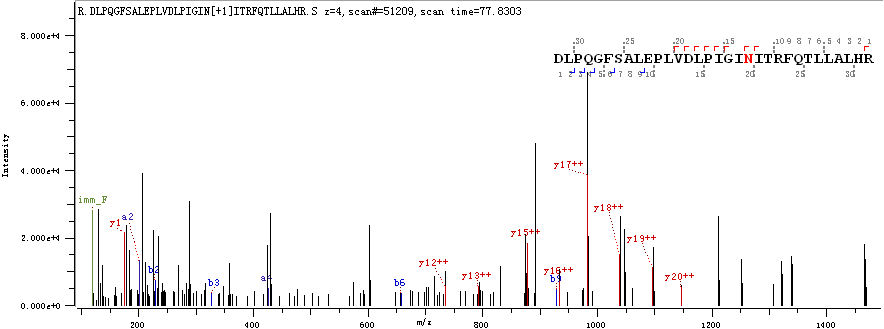

**N282**

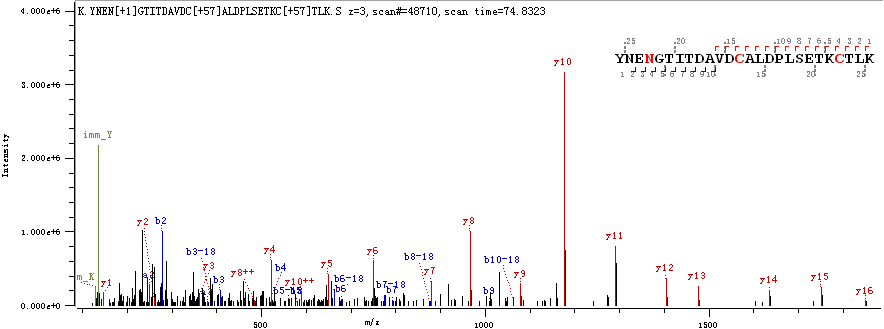

**N331**

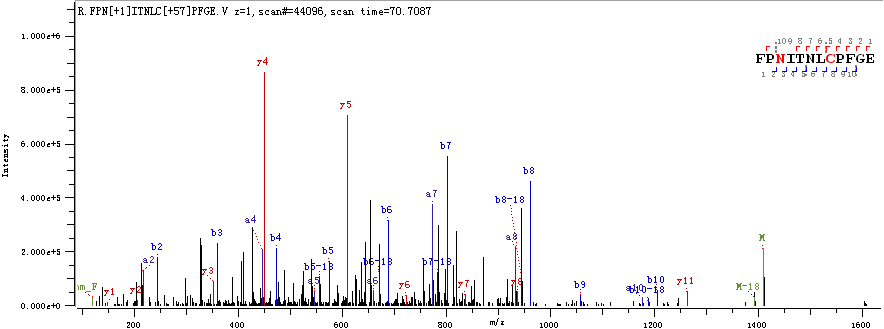

**N343**

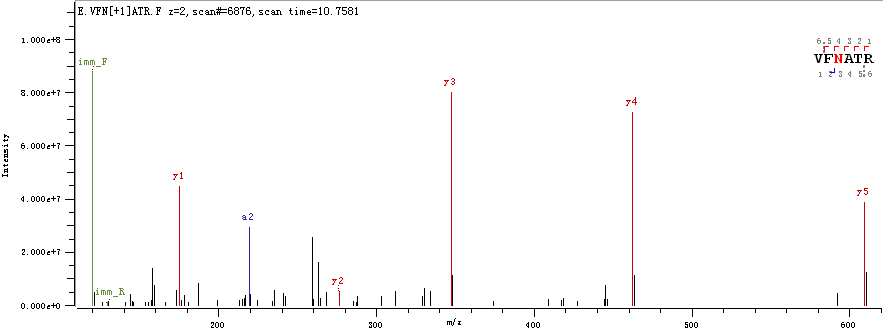

**N657**

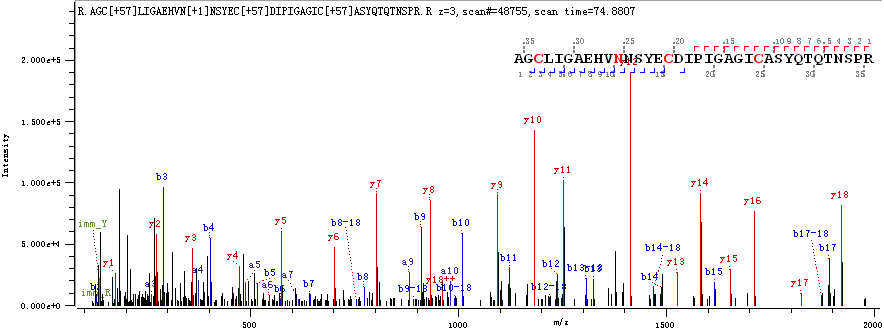

**N709**

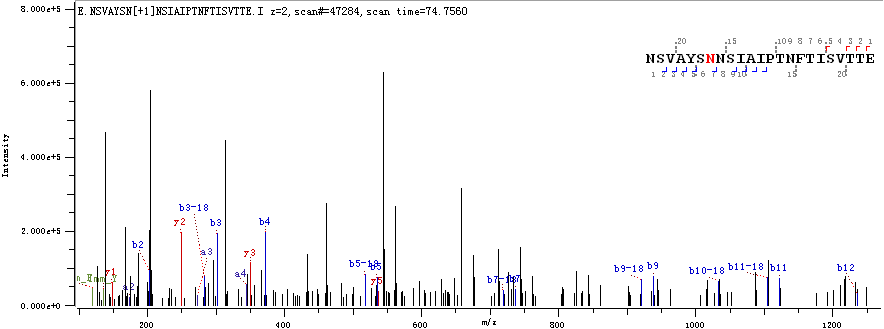

**N801**

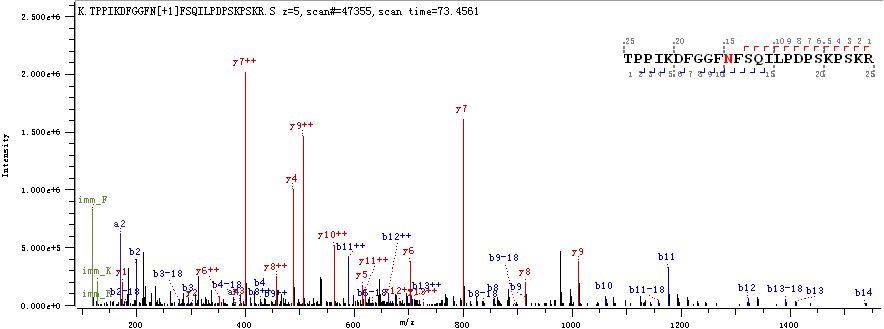

**N1074**

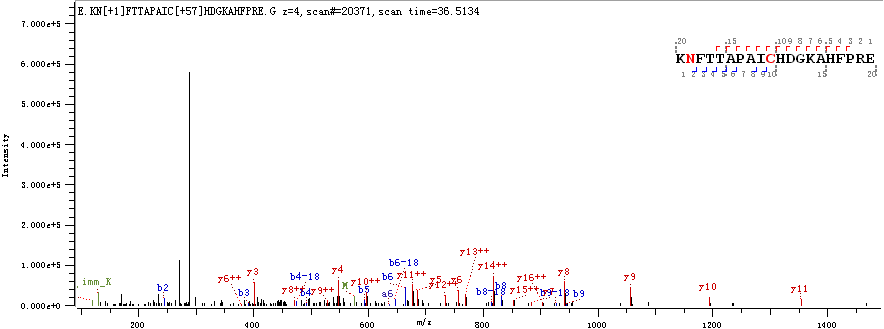

**N1098**

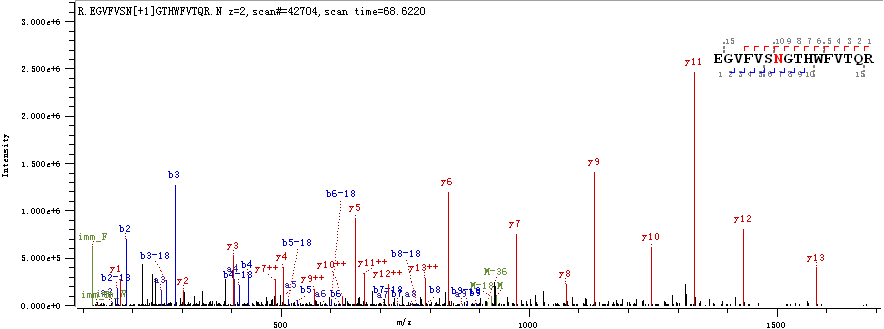

**N1158**

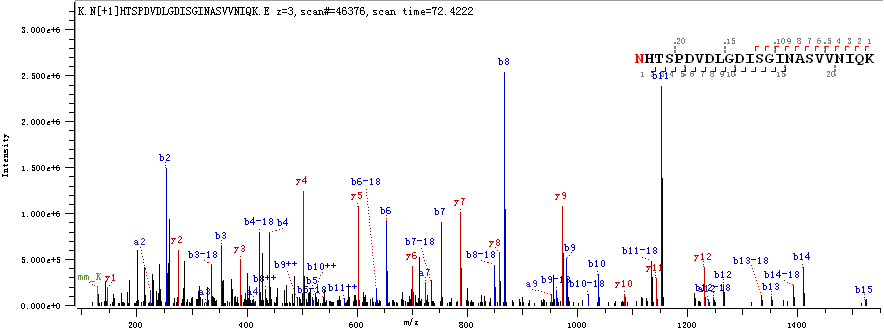

**N1173**

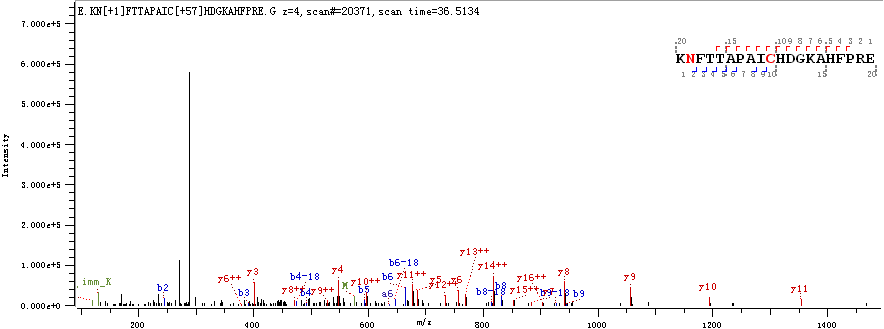

**N1194**

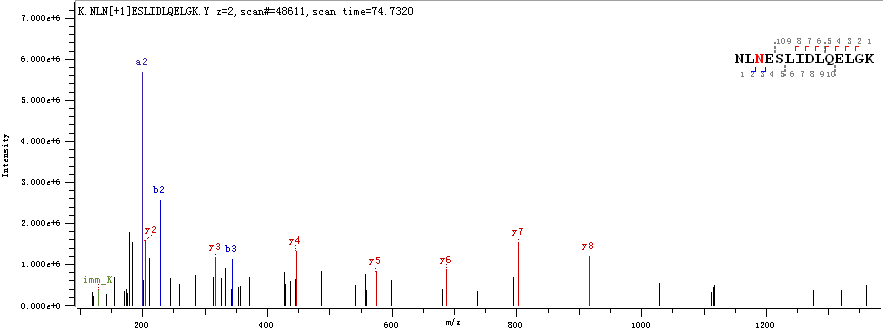

**Spectra of intact N-glycopeptides from recombinant SARS-CoV-2 spike proteins expressed in human cells**

**N17**

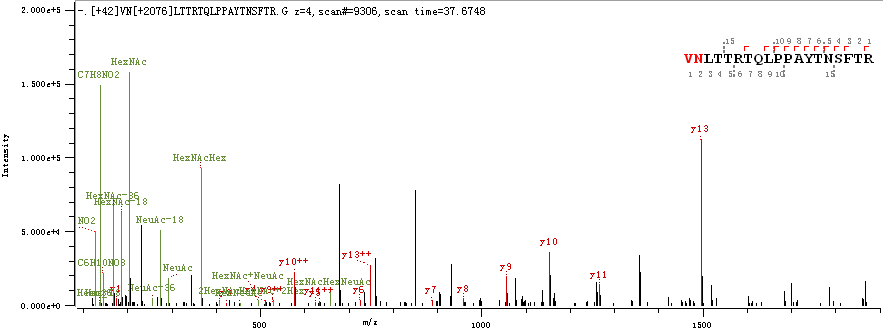

**N61 & N74**

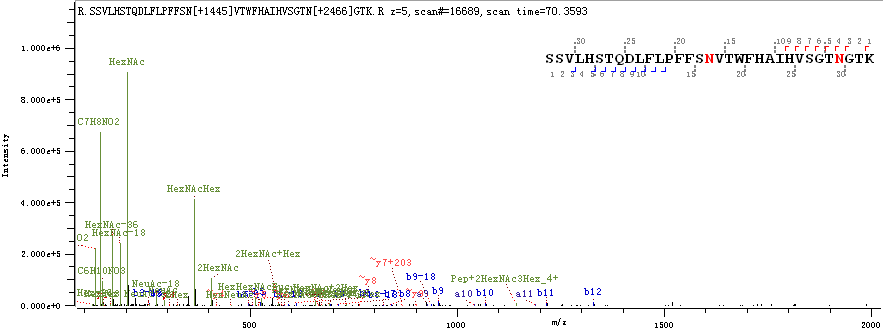

**N122**

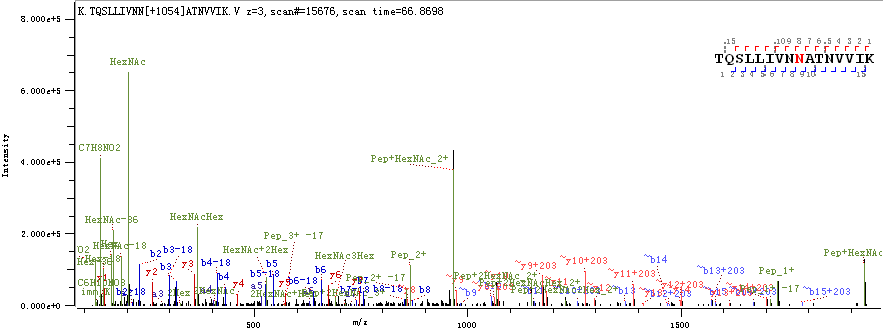

**N149**

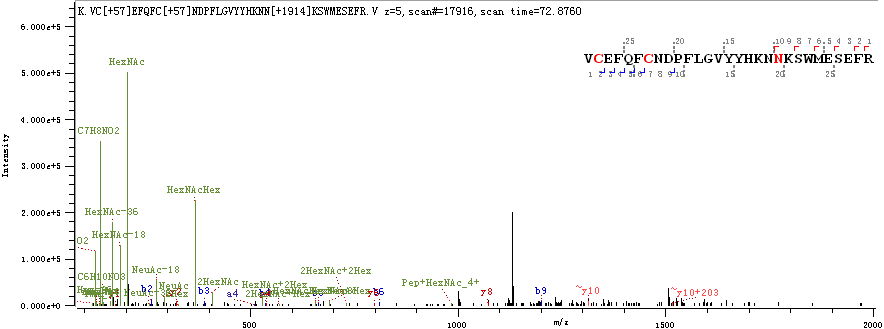

**N165**

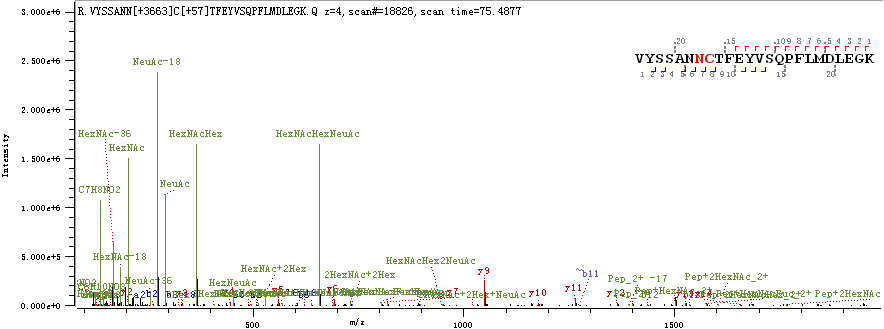

**N234**

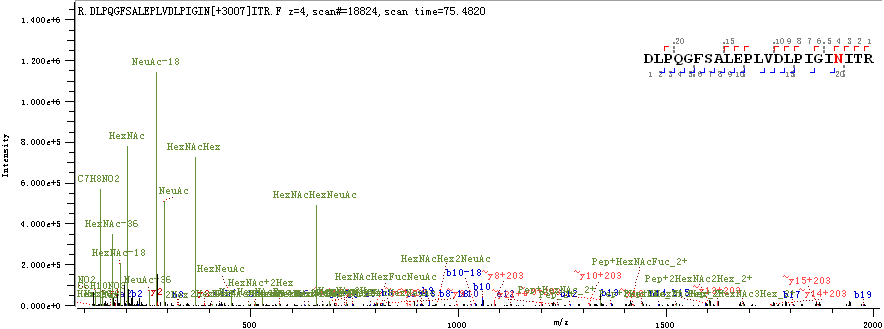

**N282**

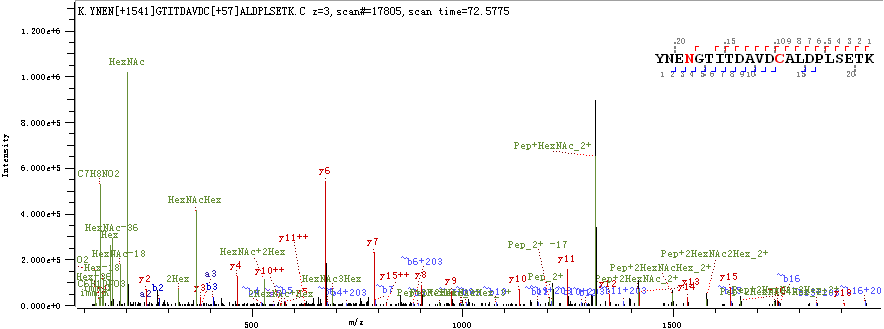

**N331 & N343**

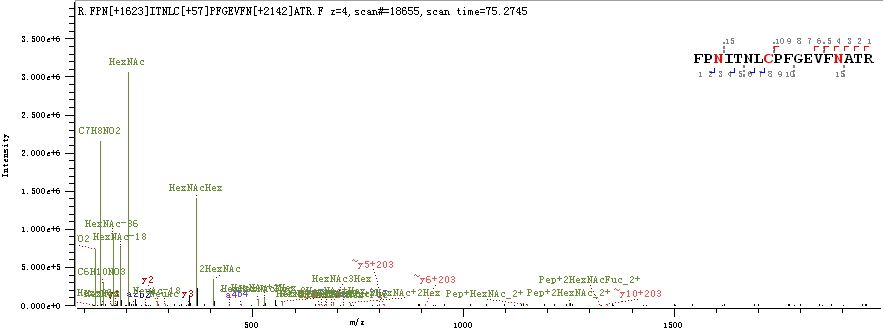

**N603 & N616**

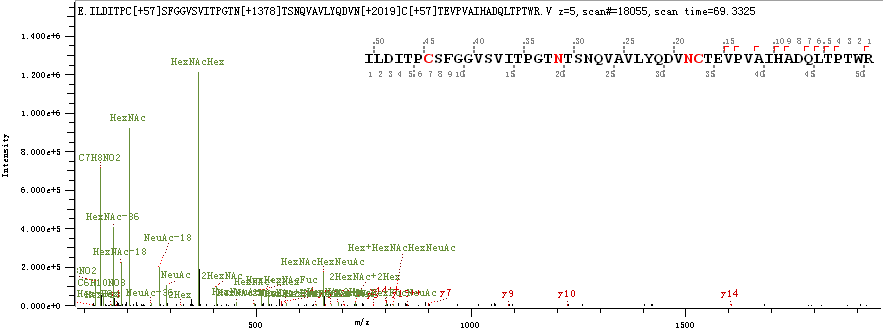

**N657**

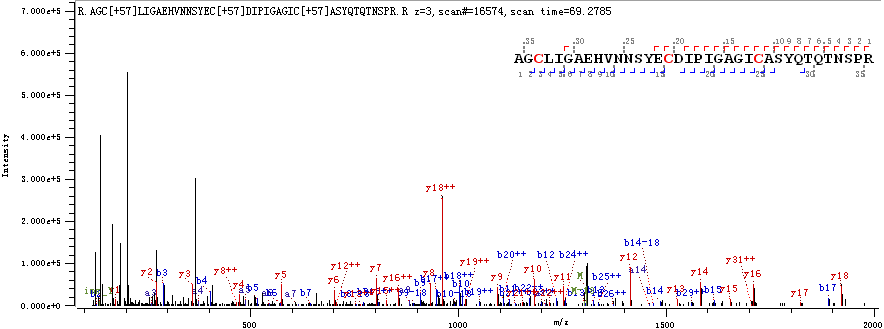

**Spectra of deglycosylated peptides from recombinant SARS-CoV-2 spike proteins expressed in human cells**

**N17**

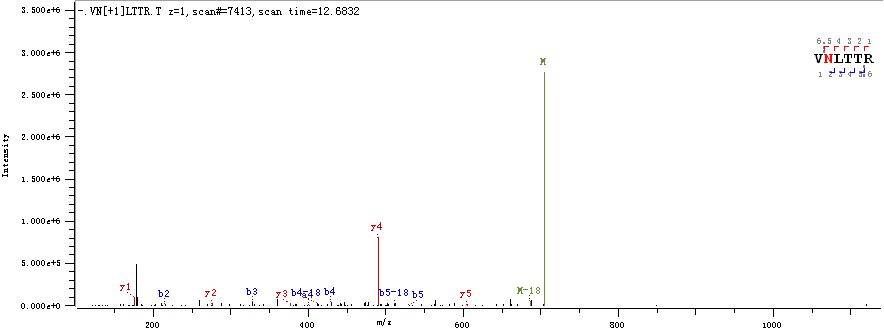

**N122**

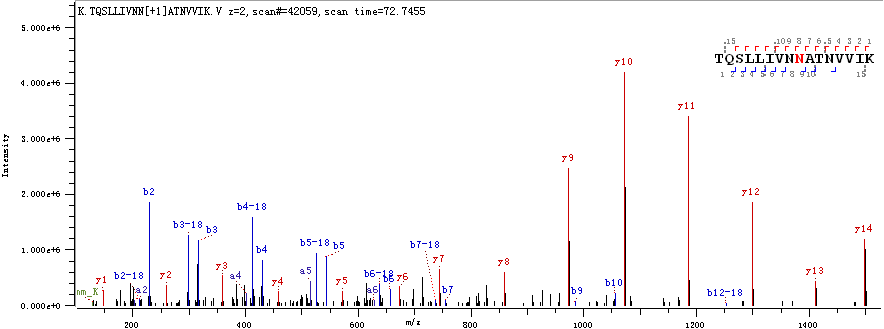

**N149**

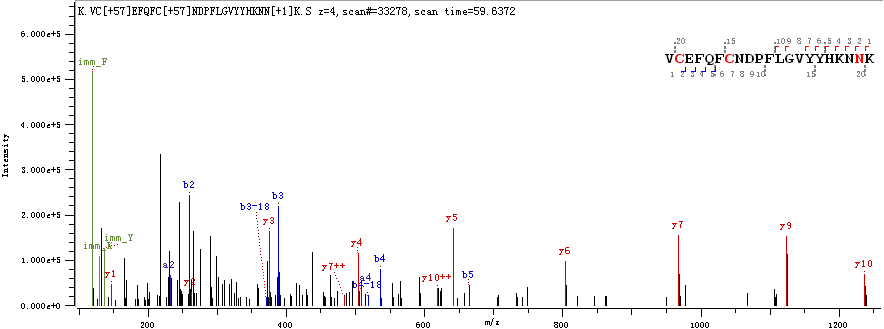

**N165**

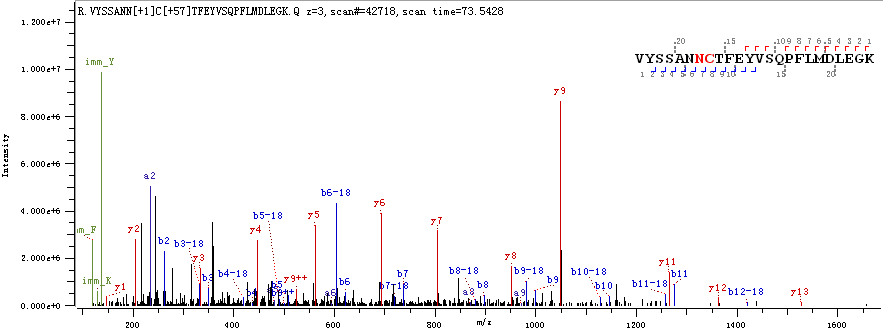

**N234**

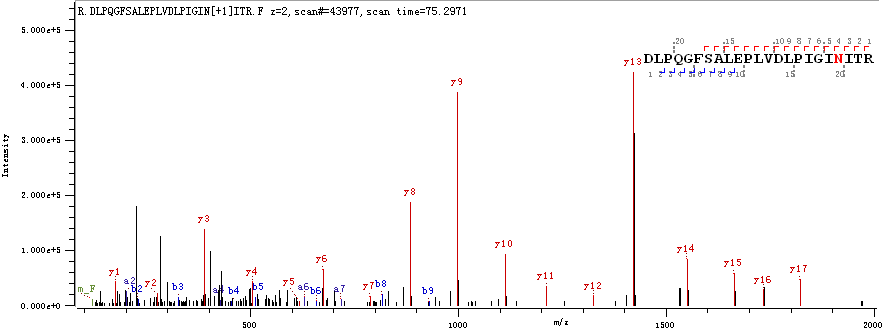

**N282**

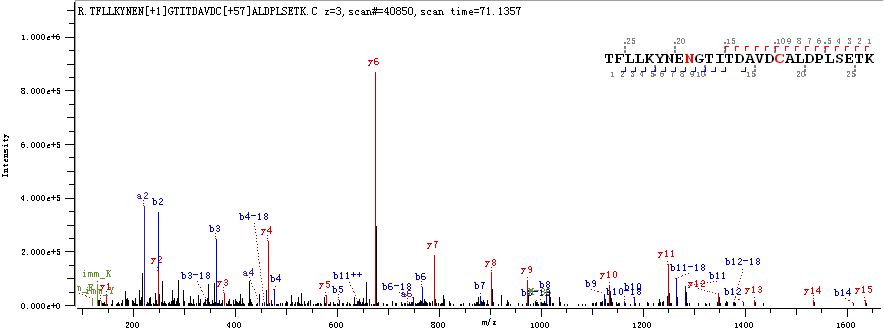

**N331**

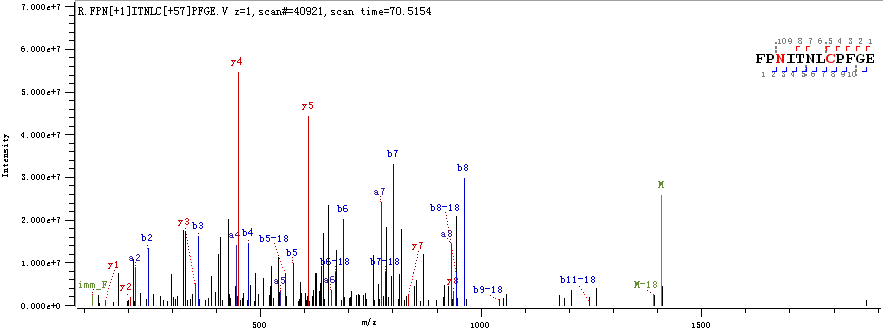

**N343**

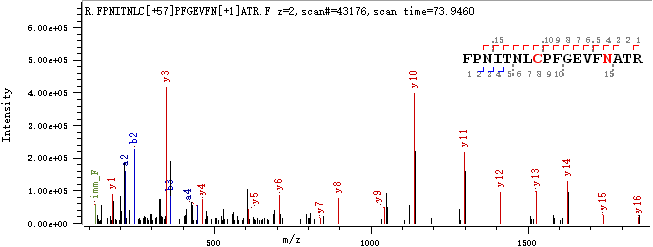

**N603**

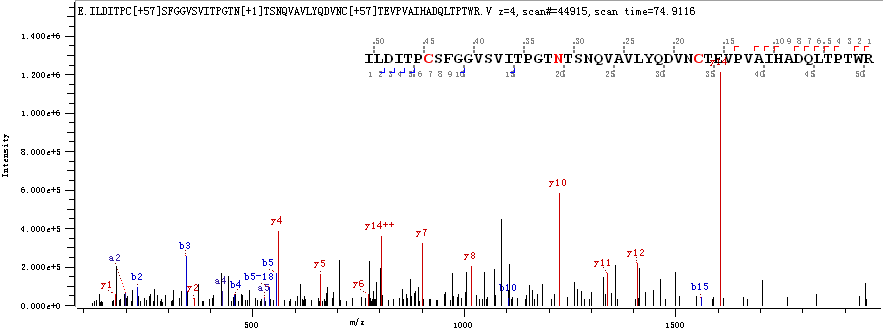

**N657**

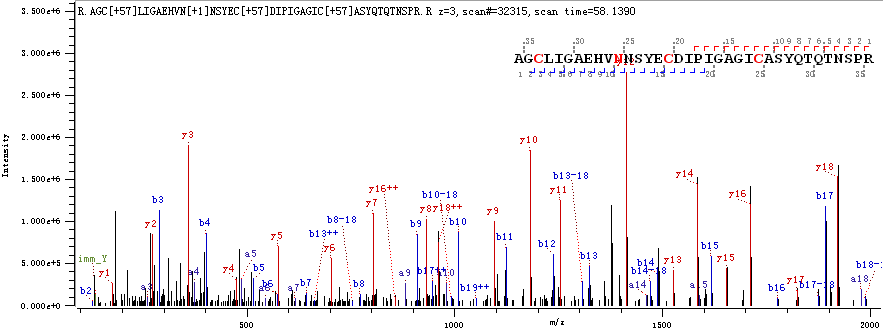

**Supplementary Figure S8.** Microheterogeneity and macroheterogeneity of the N-linked glycopeptides of the S protein.

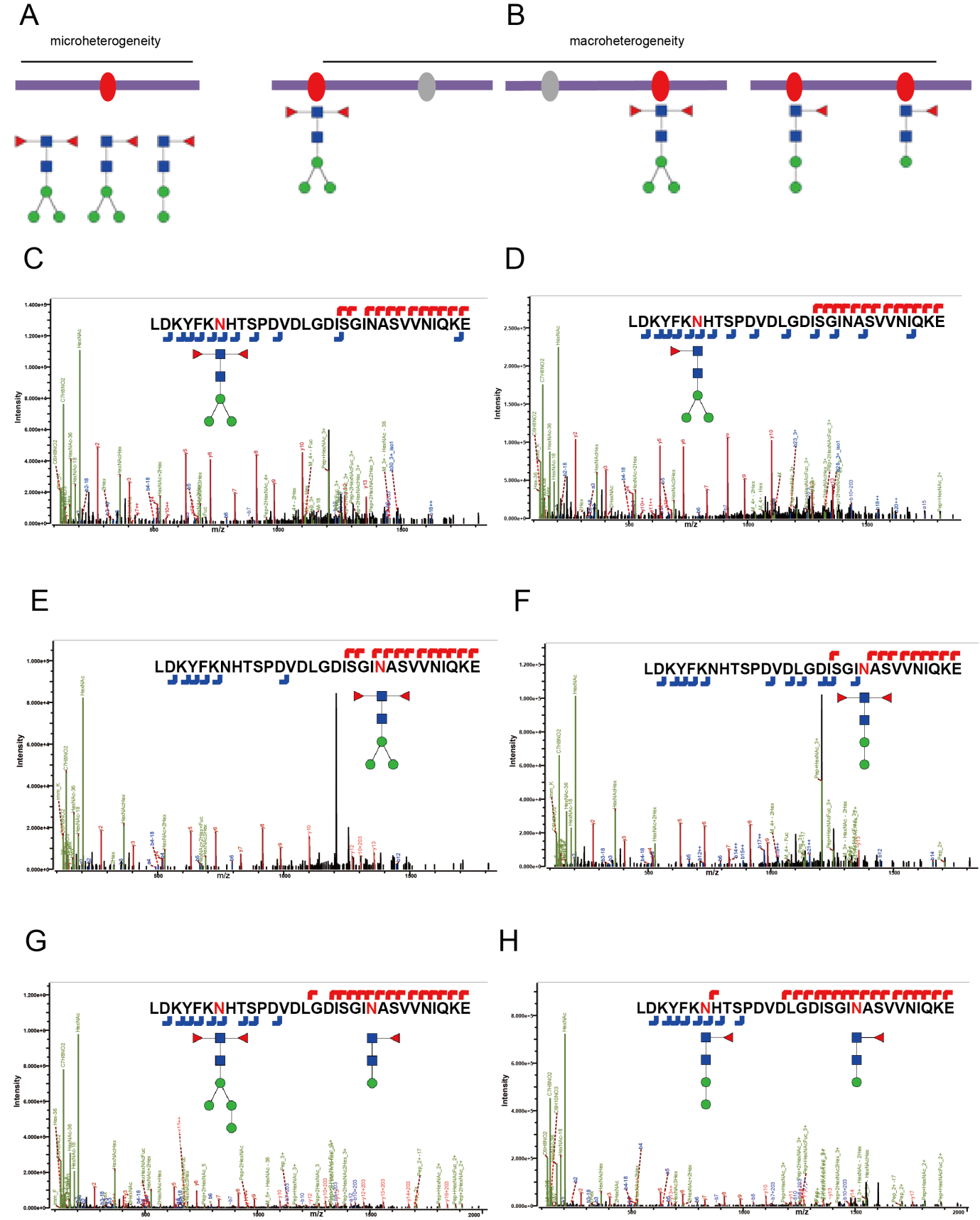

**Supplementary Figure S9.** SDS-PAGE analysis of RBDs expressed in insect and human cells.

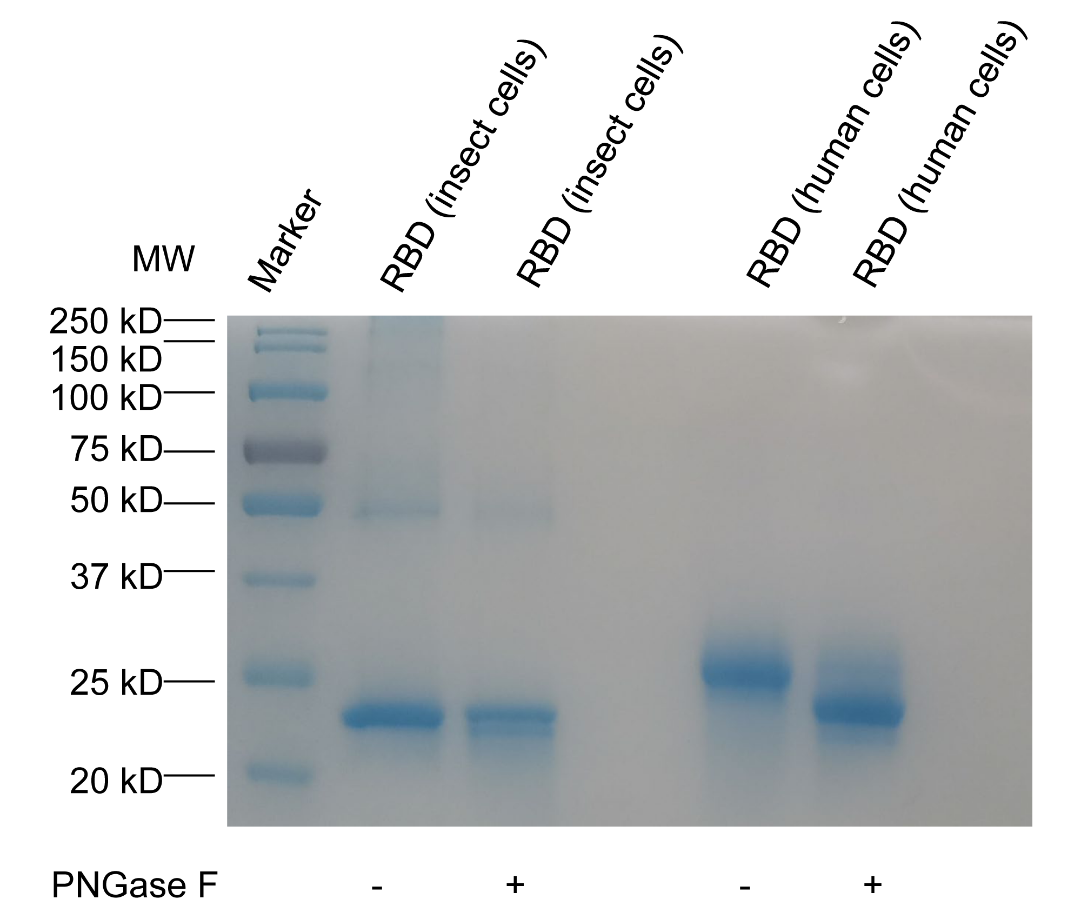
